# Climate change impacts on the abiotic degradation of acyl-homoserine lactones in the fluctuating conditions of marine biofilms

**DOI:** 10.1101/2022.01.12.476096

**Authors:** Christina C. Roggatz, Daniel R. Parsons

## Abstract

Marine biofilms are functional communities that shape habitats by providing a range of structural and functional services integral to coastal ecosystems. Impacts of climate change on biological aspects of such communities are increasingly studied, but impacts on the chemicals that mediate key interactions of biofilm organisms have largely been overlooked. Acyl-homoserine lactones (AHLs), crucial bacterial signals within biofilms, are known to degrade through pH and temperature-dependent hydrolysis. However, the impact of climate change on AHLs and thus on biofilm form and function is presently unknown. This study investigates the impact of changes in pH and temperature on the hydrolysis rate, half-life time and quantitative abundance of different AHLs on daily and seasonal timescales for current conditions and future climate change scenarios.

We established the mathematical relationships between pH, hydrolysis rates/ half-life times and temperature, which revealed that natural daily pH-driven changes within biofilms cause the greatest fluctuations in AHL concentration (up to 9-fold). Season-dependant temperature enhanced or reduced the observed daily dynamics, leading to higher winter and lower summer concentrations and caused a shift in timing of the highest and lowest AHL concentration by up to two hours. Simulated future conditions based on climate change projections caused an overall reduction of AHL degradation and led to higher AHL concentrations persisting for longer across both the daily and seasonal cycles.

This study provides valuable quantitative insights into the theoretical natural dynamics of AHL concentrations. We highlight critical knowledge gaps on the scale of abiotic daily and seasonal fluctuations affecting estuarine and coastal biofilms and on the biofilms’ buffering capacity. Detailed experimental studies of daily and seasonal dynamics of AHL concentrations and assessment of the potential implications for a suite of more complex interactions are required. Substantial fluctuations like those we show in this study, particularly with regards to concentration and timing, will likely have far reaching implications for fundamental ecosystem processes and important ecosystem services such as larval settlement and coastal sediment stabilisation.

## 1 INTRODUCTION

Climate change caused by anthropogenic carbon dioxide (CO_2_) emissions is predicted to significantly change the physical and chemical parameters of our waterbodies across Earth. Assuming a business-as-usual scenario (RCP 8.5), ocean surface pH is predicted to drop by 0.4 pH units until the end of this century, a process called ocean acidification [1]. In the same timeframe, sea surface temperature is predicted to rise by more than 4°C [1]. While the range of change is within conditions previously experienced on Earth, the rate of change is unprecedented, with severe impacts on the form and function of the environment and organisms becoming apparent.

One recently discovered effect of ocean acidification on the biospehere is that it can severely affect the molecular properties of chemical signals that mediate the interactions of marine organisms and their daily life [2]. An average change of 0.4 pH units was found to render peptides involved in crab brood-care non-functional [2] and impair hermit crabs in their ability to locate food effectively, likely due to the same reason [3]. Fishes such as sea bass and sea bream also show significant reduction in their ability to receive chemical signals in reduced pH conditions [4, 5]. When a chemical signal is transported from the source or sender to the receiving organisms, it is subject to the environmental conditions within which it is transported and will therefore inevitably be affected by the surroundings. Climate driven changes to these surroundings will thus likely have a suite of poorly understood impacts on signal used for chemical communications between organisms.

Biofilms are ubiquitously distributed worldwide within estuarine and coastal settings, providing a range of structural and functional services that are integral to coastal ecosystems and morphological stability [6, 7]. N-acyl-homoserine lactones (AHLs) are key signalling molecules used by bacteria in cell-cell communication and play a crucial role in biofilm formation and the production of extracellular polymeric substances (EPS) [8]. The importance of these signals in marine, estuarine and coastal microbial mats and biofilms, however, only came into focus in the past 20 years. In 2002, the production of AHLs within Roseobacter and Marinobacter strains isolated from marine snow was reported for the first time [9]. Since then a variety of AHL producing microorganisms, mainly gram-negative bacteria, have been isolated from marine biofilms [10, 11] (and references therein). Due to the very low concentration of AHLs in environmental samples, only few studies managed to identify and quantify these compounds directly. Decho *et al*. extracted, identified and quantified nine different AHLs from stromatolite microbial mats, of which C6-, C8-and C10-HSL were particularly abundant [12]. Tait and co-workers were able to extract AHLs from rock-pool pebble-biofilms averaging a concentration of approximately 600 pmol cm^−2^ and found C8- and C10-HSL to dominate [13]. More recently, AHLs were also quantified in intertidal marine sediments with C8-, C10- and C12-HSL dominating the profile [14]. Besides their presence in marine bacterial biofilms, where AHLs mediate the bacteria-bacteria interactions via quorum sensing, it was shown that AHLs are further involved in a number of cross-kingdom interactions [15]. C10-HSL, its 3-oxo and 3-OH forms, have been found to mediate interactions between benthic diatoms and bacteria [16] while a range of AHLs from C6-HSL to C14-HSL and their hydroxyl- and oxo-forms were found to act as attractants for larvae of macro algae [17, 18] and biofouling or bioturbating fauna [19] (and references therein) (see Fig. 1A for an overview of AHL-mediated interactions).

**Figure 1:**
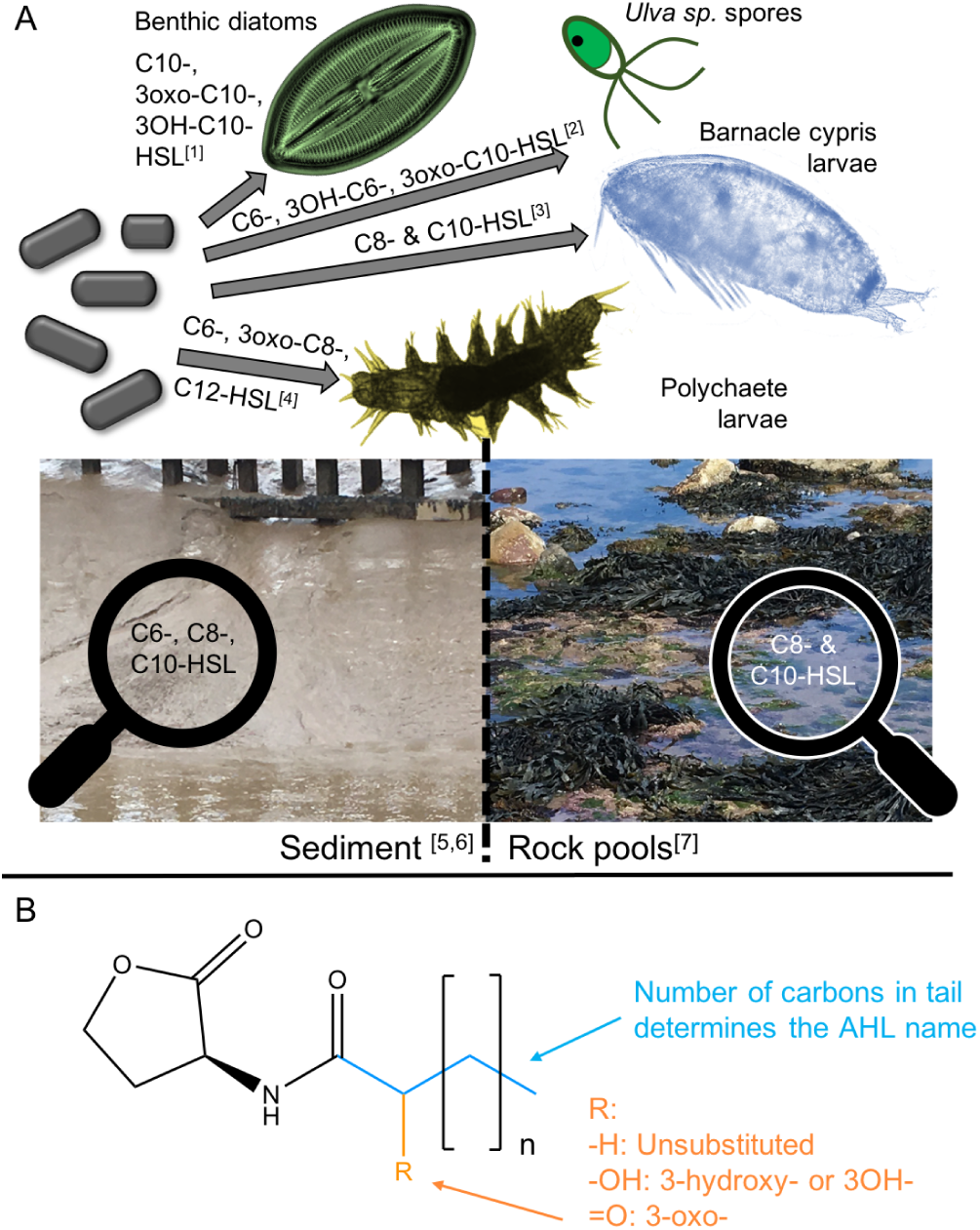
Overview of AHL-mediated interactions, their presence in different environments and their basic chemical structure. Panel A gives an overview of bacterial AHLs that are known to mediate interactions with selected algae and invertebrates, and indicates important AHLs for different marine habitats. Panel B shows the basic structure of AHLs and explains the nomenclature with the number of carbons in the blue tail giving the name and the substitution at R in orange determining the AHL class. References: [1] Yang *et al*. [16], [2] Joint *et al*. [18], [3] Tait & Havenhand [19] and [4] Huang *et al*. [20], [5] Decho *et al*. [12], [6] Stock *et al*. [14] and [7] Tait *et al*. [13].

All N-acyl-homoserine lactones follow a common structure consisting of a homoserine lactone ring, which is N-acylated with a fatty acyl group at the *α*-position [21]. The fatty acid group can be of variable acyl chain lengths (usually 4 to 18 carbons), saturation levels and oxidation states, belonging to either the N-acyl, N-(3-oxoacyl) or N-(3-hydroxyacyl) class (see Fig. 1B). [21] In this study, names of AHLs are abbreviated in the common way by using Cx or Cx-HSL (interchangeably) with x = number of carbons. Importantly, this structure makes AHLs susceptible to change with pH. In fact, only AHLs with chain lengths of C ≥ 4 persist long enough to convey a signal [22, 23]. The pH-dependent, base-catalysed hydrolysis of the lactone ring transforms the AHL into the corresponding N-acylhomoserine, which no longer functions as a chemical signal [22, 24]. This reaction is further accelerated by increasing temperatures [22]. AHLs are therefore assumed to be short-lived signalling cues, especially those with short side chains of six carbons or less, which degrade quickly in marine environments with pH > 7. [25] Degradation of AHLs in seawater was established experimentally by Tait *et al*. [17] and Hmelo & Van Mooy [23] for AHLs with a range of chain lengths and substitutions. Decho and coworkers went one step further and measured the pH profile within microbial mats under natural conditions and then experimentally quantified the half-life time of some AHLs in different pH conditions during laboratory studies. They established a significant degradation of the shorter chain AHLs in the laboratory and in the natural microbial mat during daytime in the field, and subsequently linked their observations to the significant daily pH fluctuations they observed within the biofilm. [12] However, despite numerous publications highlighting and studying the influence of environmental physical parameters on AHL signalling in general, the impact of naturally fluctuating abiotic conditions within and in the surrounding of biofilms remains undetermined, as highlighted by Decho & Gutierrez [26] as well as Hmelo [25] in recent reviews. The impacts of seasonal variations, and/ or climate change scenarios, have not been addressed to date.

This study therefore investigates the impact of changes in pH and temperature on the quantitative abundance of different AHLs for daily and seasonal conditions in the context of current and future climate change scenarios. First, the mathematical relationships between pH and each specific AHL hydrolysis rate *k* and half-life time *t*_1*/*2_ as well as the influence of temperature on *k* and *t*_1*/*2_ are established. Then the change in AHL hydrolysis rate, half-life time and relative concentration is calculated for daily fluctuations within the biofilm, for seasonal variations of conditions and for average ocean conditions based on climate change projections. Finally, the scale of influence through natural fluctuations and changes due to climate change are compared and the implications for interactions mediated through AHLs are discussed in terms of the ecosystem services and stability of coastal and estuarine systems.

## 2 MATERIALS & METHODS

### 2.1 AHL hydrolysis kinetics and pH

The degradation of AHL due to hydrolysis in water (also called lactonolysis) follows a pseudo first-order reaction. For the neutral and alkaline hydrolysis of interest in the context of this study, the reaction follows a B_AC_2 mechanism as described by Goméz-Bombarelli and colleagues [27]. The reaction can be described as

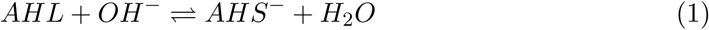

where AHL stands for N-acyl-homoserine lactone and AHS for the corresponding N-acyl homoserine. With the reaction taking place in water, the hydrolysis rate *k* at any given condition can be calculated as:

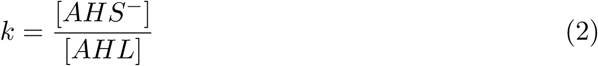

following the pseudo-first order as shown by Ziegler *et al*. [28]. The hydrolysis rate *k* can further be converted into half-life time *t*_1*/*2_ using

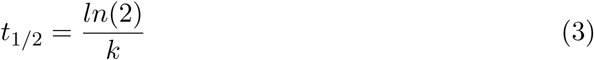

However, the hydrolysis rate and half-life time of AHLs are molecule-specific and further dependent on pH, temperature and the length of their alkyl-chain [22].

#### 2.1.1 Dependence of the hydrolysis rate *k* on pH

As can be seen from eqn. (1), the concentration of hydroxide anions ([OH^−^]) and therefore pH plays a central part in the hydrolysis of AHLs. Limited [OH^−^] will slow hydrolysis down while higher concentrations or even excess of [OH^−^] will accelerate the ring-opening reaction. In order to obtain a general mathematical relationship for the dependency of *k* on pH, we formulate the pH-dependent rate *k*_*pH*_ based on eqn. (1) as

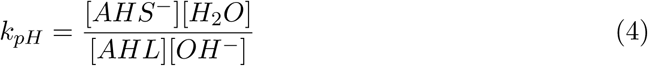

The concentration of hydroxide anions is liked to pH through

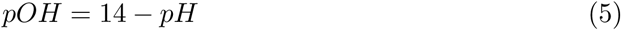

and

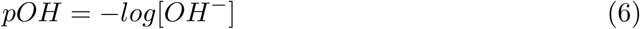

so

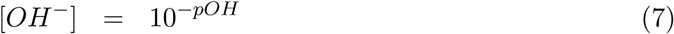

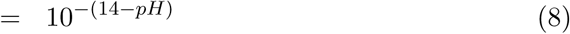

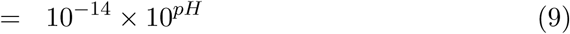

Considering in this context [H_2_O] = 10^−14^ and substituting [H_2_O] and [OH^−^] into eqn. (4) yields

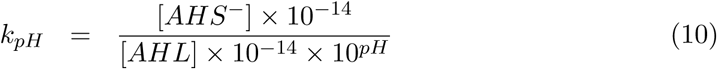

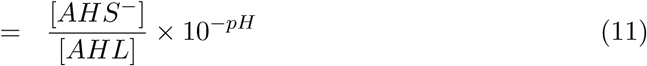

which can then be expressed as a linear relationship by multiplying with the negative decadic logarithm

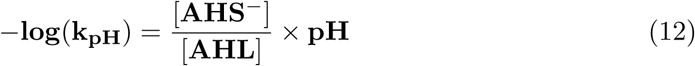

to describe the link between the AHL/AHS ratio and pH.

In order to establish the AHL-specific coefficients for this equation, the data published by Ziegler *et al*. [28] has been used, who measured the pH-specific hydrolysis rates of C4, C6, C8, C6-oxo and C8-oxo by ^1^H NMR spectroscopy in D_2_O at pH 7.0, 7.9, 9.2 and 9.5 at room temperature (22°C). The rates were plotted as negative decadic logarithm versus the pH in IGOR pro (v6.37) and a linear least-square fit function was obtained. The slope of the fit function represents the 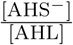 coefficient, which can subsequently be used to calculate the AHL-specific k_pH_ at any given pH.

The same analysis was performed for data obtained by Decho *et al*. [12], who published the half-life time of C6, C8, C10, C12 and C14 at pH 6.18, 7.2, 8.2, 8.7 and 9.55 recorded at 26°C. The *t*_1*/*2_ data was converted into *k* using eqn. (3) and analysed as described above.

#### 2.1.2 Dependence of the half-life time *t*_1*/*2_ on pH

For the dependence of the AHL-specific half-life time *t*_1*/*2_ on pH, a similar relationship as for the hydrolysis rate can be established by substituting eqn. (3) into eqn. (11).

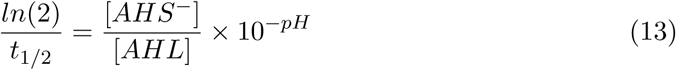

and rearranging to

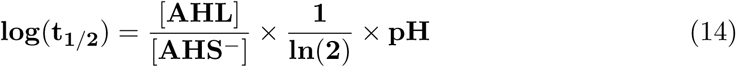

Like for *k*_*pH*_, the data sets by Ziegler *et al*. [28] and Decho *et al*. [12] were used. For consistency, all times were transformed to minutes.

### 2.2 AHL hydrolysis kinetics and temperature (T)

The impact of temperature on biological and chemical processes is often expressed through a temperature coefficient (mostly for steps of 10°C, hence Q_10_). It assumes, that the reaction rate (or in this study the hydrolysis rate *k*) depends exponentially on the temperature T. For a 1°C temperature change, Q_1_ can be expressed as

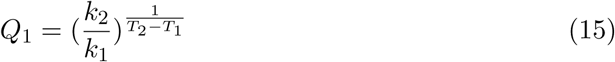

where *k*_1_ and *k*_2_ are the hydrolysis rates at two different temperatures, *T*_1_ and *T*_2_, respectively. The temperature step of 1°C is represented by the 1 in the exponential fraction.

Based on the investigations of Yates *et al*. [22], who report the hydrolysis rates for C4, C6, C8 and C6-oxo at a stable pH and 22 or 37°C, the temperature coefficient per 1°C was calculated using eqn. (15).

#### 2.2.1 Dependence of the hydrolysis rate *k* on T

To account for the impact of temperature on the hydrolysis rate, the rates obtained for C4, C6, C8 and C6-oxo at any given pH were shifted to the desired temperature by multiplication with the coefficient according to the temperature difference. For example, to shift *k* of C6-HSL obtained at pH 8.2 and 26°C to a more relevant temperature of 16 or 20°C, *k* was multiplied by Q_−10_ or Q_−6_, respectively.

#### 2.2.2 Dependence of the half-life time *t*_1*/*2_ on T

For the influence of T on *t*_1*/*2_, the same approach was taken based on the data of Yates *et al*. [22]. The reported relative hydrolysis rates were transformed into half-life times prior to the calculation of the *t*_1*/*2_-influencing coefficient.

### 2.3 Quantification of change to *k, t*_1*/*2_ and [AHL] for natural conditions and climate change scenarios

#### 2.3.1 Definition of relevant natural pH and temperature ranges

Physical aquatic parameters, such as temperature and pH, fluctuate considerably in natural environments like estuaries [29, 30]. Defining relevant pH and T ranges is therefore crucial. Although AHL-producing bacteria in biofilms are assumed to be well-buffered from the surroundings [31], pH conditions within the biofilm can vary considerably due to the presence of photosynthetic co-inhabitors such as diatoms [12] (and references therein). Decho *et al*. [12] showed that within the first millimetres of a marine biofilm, pH can fluctuate between an acidic pH 6.8 at night and alkaline pH of 9.4 during the day at a stable external pH. This pattern can be translated into a sinus function that represents pH over the course of a day:

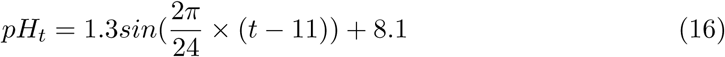

with *t* specifying the hour of the day out of 24.

#### 2.3.2 Natural conditions in the Humber estuary

Abiotic water parameters, such as pH and temperature, have been measured frequently over the past years within and surrounding the Humber estuary (UK). For this study the dataset for pH and temperature measured at Spurnpoint, Saltend Jetty and Albert Dock from 1995 to 2005 was used (available upon request from corr. author, will be made available for publication). Data was pooled and plotted with respect to the day within the year it was obtained, before being analysed for apparent fluctuations. pH showed some variation (pH 7.78 ± 0.23), but no clear temporal or spatial pattern is evident. Temperature data (11.15 ± 4.86 °C) showed a clear seasonal pattern and was subsequently fitted with a sinus function in IGORpro (v6.3).

#### 2.3.3 Relevant climate change scenarios

Based on the latest IPCC report, global average surface ocean pH is currently assumed as pH 8.1 and predicted to drop by 0.4 units to pH 7.7 by the end of this century [1]. Global average surface ocean temperature is currently at 16°C [32] and predicted to rise to 20°C by 2100 [1]. Assuming any average changes predicted with climate change would translate to the biofilm environment unaltered (i.e. cause a baseline shift), the natural pH conditions in the biofilm by the year 2100 could be shifted to range from 6.4 to 9.0 and temperature could be increased by up to 4°C.

#### 2.3.4 Calculation of scenario-specific *k, t*_1*/*2_ and relative [AHL]

For this part only the effects for C6 and C8 were evaluated as data for these two AHLs with regards to pH and temperature influences was most reliable. To account for potential experimental uncertainties, the hydrolysis rate and half-life time data for C6 and C8 obtained by Decho *et al*. [12] and Ziegler *et al*. [28] was combined after adjusting the NMR-based data of Ziegler and co-workers to the temperature used for Decho and co-workers’ experiments (based on the temperature coefficients obtained through Yates *et al*. [22]) and accounting for the kinetic isotope effect (KIE) of D_2_O compared to water 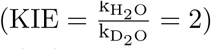 by multiplying Ziegler’s hydrolysis rates by 0.5 and the respective half-life times by 2 [28]. The combined dataset was then plotted against pH and subjected to to the analyses described above to obtain the respective linear correlation coefficients.

Then, for each specifically defined condition (e.g. each datapoint of the seasonal Humber dataset or each combination of average climate change conditions), the pH and T-dependent hydrolysis rate *k* and the respective half-live time *t*_1*/*2_ were calculated based on eqn. (12) and (14) and the corresponding temperature coefficients based on (15). Differences between maximum, average and minimum of fluctuating conditions or between current and future average conditions were calculated and expressed in plain numbers as well as % (relative to average or current conditions). Monthly averages for seasonal variations were calculated and expressed as ± standard error of mean (SEM). Seasonal trends for hydrolysis rate and half-life time across the year were analysed by fitting a sine function (IGORpro v6.3) based on the observations that temperature the is most influencing factor.

For each climate change scenario the AHL concentration over time (minutes) was calculated based on a classic exponential decay equation and assuming an AHL start concentration of 1, so

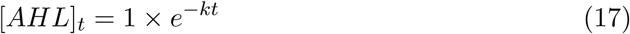

using the respective hydrolysis rate *k* adjusted for pH and T of the scenario in question. For the daily periodical fluctuations, a constant hourly production of [AHL] = 1 was assumed and summing up all produced and from decay remaining hour-specific AHL concentrations (calculated based on eqn. (16)) yielded the overall relative [AHL] for each hour of the day.

## 3 RESULTS

### 3.1 Numerical pH-dependence of hydrolysis rate and half-live time

For the investigated pH range between 6.0 and 10.0, there was a clear linear impact of pH on the hydrolysis rate when plotted at negative log-scale (Fig. 2). The same could be observed for half-life time (Fig. S1). With increasing pH, −*log*(*k*) decreased, corresponding to an increase of the hydrolysis rate *k*. At lower pH conditions the hydrolysis rate is slower.

**Figure 2:**
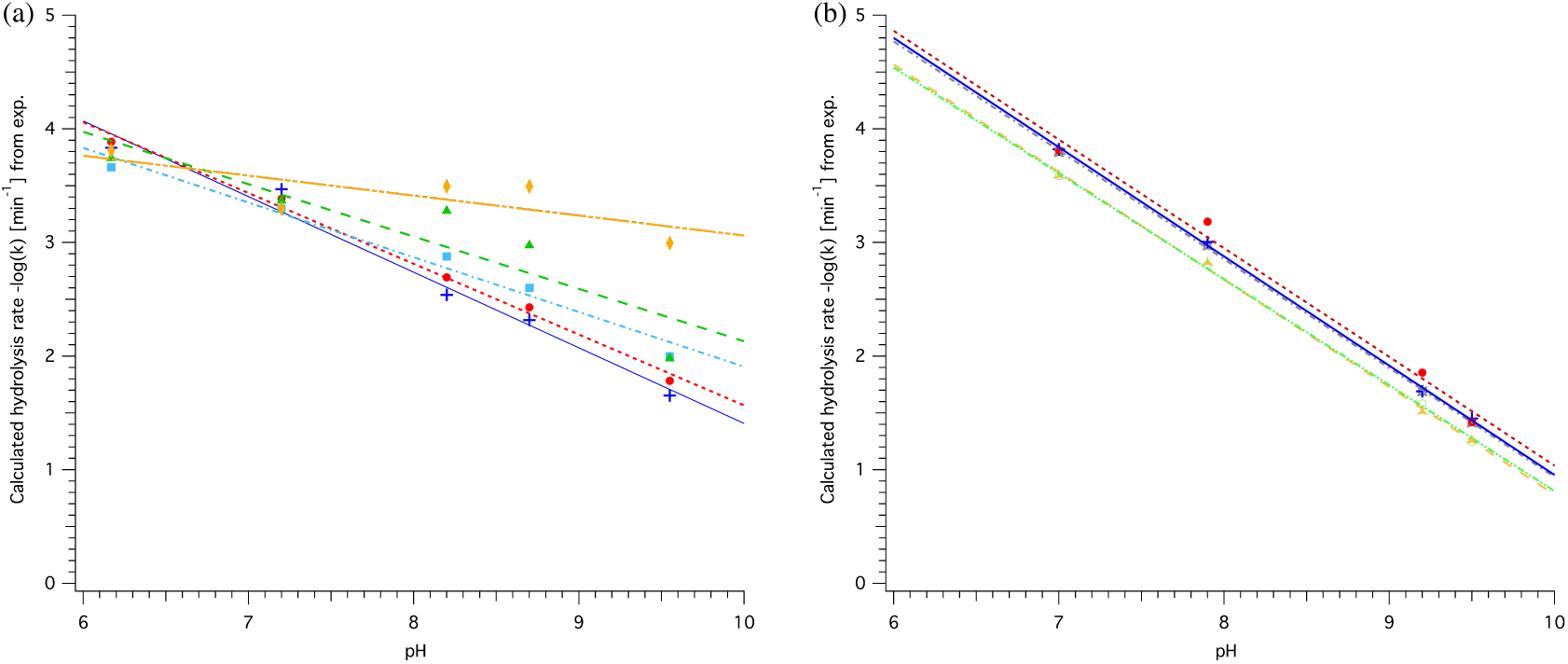
pH-dependence of hydrolysis rate *k* for different AHLs. (a) Based on data published by Decho *et al*. [12] for C6 (blue, cross), C8 (red, circles), C10 (light blue, squares), C12 (green, triangles) and C14 (orange, diamonds). (b) Based on data published by Ziegler *et al*. [28] for C4 (grey, star), C6 (blue, cross), C8 (red, circles), C6-oxo (orange, triangle) and C8-oxo (light green, open circles). Coefficients for linear least-square fit following the equation −*log*(*k*) = *a* × *pH* + *C* are detailed in Table 1.

The steeper slope observed for AHLs with shorter acyl-chain length based on the data by Decho *et al*. [12] indicates a greater impact of pH on the hydrolysis rate than for AHLs with longer acyl-chains with more than 8 carbons (see also slope coefficient *a* in Table 1). For C10-HSL the slope was 0.14 lower than for C8-HSL, a significant reduction by more than 22%. In addition, an overall faster abiotic hydrolysis rate for shorter chain AHLs is reflected by the calculated *k* at pH 8.0 in Table 1. For AHLs with 8 carbons or less, the parameters are similar, suggesting a similar impact of pH and similar rate of abiotic hydrolysis (see (a) and (b) in Fig. 2). AHLs with a 3-oxo-group in the side chain had a faster hydrolysis rate at each point across the pH range, but were equally affected by pH (similar slope). It has to be noted that the least-square linear fit obtained for C6, C8 and C10 data by Decho *et al*. [12] as well as for all data by Ziegler *et al*. [28] was very good (R^2^ > 0.95), while the linear regression obtained for C12 and C14 was not as good or even poor and the subsequently calculated data, such as *k* at any pH, should be interpreted with caution. It also has to be taken into consideration that rates obtained based on either dataset are only representative for the respective conditions. While Decho *et al*. [12] obtained their rates in water at 26°C based on subsequent GC-MS analyses in 0.5h steps, Ziegler *et al*. [28] performed NMR experiments at 22°C in D_2_O and determined *k* based on sampling in steps of 4min. Hence the obtained parameters for C6 and C8 by both groups are not directly comparable without temperature and solvent adjustment.

**Table 1:** Coefficients (± SD) of the relationship between hydrolysis rate *k* and pH. expressed as linear equation of the form -log(*k*) = *a* x pH + *C*, valid for the pH range from 6.0 to 10.0. R^2^ expresses the goodness of fit of the linear regression. *k* at pH 8.0 calculated based on fit equation and expressed as rate per minute.

| AHL | $a$ | $C$ | $R^2$ | $k$ at pH 8.0<br>($\times 10^{-3}$) [$\text{min}^{-1}$] | Source data |
| --- | --- | --- | --- | --- | --- |
| C6 | -0.67 ± 0.06 | 8.1 ± 0.4 | 0.980 | 1.8 | Decho *et al.*<br>(2009)* |
| C8 | -0.62 ± 0.03 | 7.8 ± 0.2 | 0.994 | 1.4 | |
| C10 | -0.48 ± 0.04 | 6.7 ± 0.4 | 0.975 | 1.4 | |
| C12 | -0.5 ± 0.1 | 6.7 ± 0.9 | 0.822 | 2.0 | |
| C14 | -0.18 ± 0.08 | 4.9 ± 0.7 | 0.592 | 0.3 | |
| C4 | -0.96 ± 0.01 | 10.5 ± 0.1 | 0.999 | 1.4 | Ziegler *et al.*<br>(2019)** |
| C6 | -0.96 ± 0.02 | 10.6 ± 0.2 | 0.999 | 1.2 | |
| C8 | -0.96 ± 0.07 | 10.6 ± 0.6 | 0.989 | 1.2 | |
| C6-oxo | -0.95 ± 0.02 | 10.2 ± 0.2 | 0.999 | 2.5 | |
| C8-oxo | -0.93 ± 0.02 | 10.2 ± 0.2 | 0.999 | 2.2 | |

\* in H<sub>2</sub>O at 26°C, \*\* in D<sub>2</sub>O at 22°C

For the half-life time the same linear impact of pH could be observed when plotted at positive log-scale (Fig. S1). Increased pH results in a shorter half-life time, which is also illustrated in Table 2. pH has a stronger effect on short acyl-chain AHLs, which also have an overall shorter half-life, for example comparing C6, C8 and C10 at pH 8.0. As half-life time and hydrolysis rate can be simply inter-converted using equation (3), the observed trends are thus essentially the same.

**Table 2:** Coefficients (± SD) of the relationship between half-life time *t* _*1/2*_ and pH. expressed as linear equation of the form log(*t*_*1/2*_) = *b* x pH + *D*, valid for the pH range from 6.0 to 10.0. R^2^ expresses the goodness of fit of the linear regression.

| AHL | $b$ | $D$ | $R^2$ | $t_{1/2}$ at pH 8.0<br>[min] | Source data |
| --- | --- | --- | --- | --- | --- |
| C6 | -0.67 ± 0.06 | 7.9 ± 0.4 | 0.980 | 347 | Decho *et al.*<br>(2009)* |
| C8 | -0.62 ± 0.03 | 7.6 ± 0.2 | 0.994 | 437 | |
| C10 | -0.48 ± 0.04 | 6.6 ± 0.4 | 0.975 | 575 | |
| C12 | -0.5 ± 0.1 | 6.6 ± 0.9 | 0.822 | 398 | |
| C14 | -0.18 ± 0.08 | 4.7 ± 0.7 | 0.592 | 1820 | |
| C4 | -0.97 ± 0.02 | 10.5 ± 0.1 | 0.999 | 550 | Ziegler *et al.*<br>(2019)** |
| C6 | -0.97 ± 0.02 | 10.5 ± 0.2 | 0.999 | 550 | |
| C8 | -0.96 ± 0.07 | 10.4 ± 0.6 | 0.989 | 525 | |
| C6-oxo | -0.95 ± 0.02 | 10.1 ± 0.2 | 0.999 | 316 | |
| C8-oxo | -0.93 ± 0.02 | 10.0 ± 0.2 | 0.999 | 302 | |

\* in H<sub>2</sub>O at 26°C, \*\* in D<sub>2</sub>O at 22°C

The impact of temperature on the hydrolysis of C4, C6, C8 and C6-oxo has already been established by Yates *et al*. [22]. Based on their data, the general temperature coefficients shown in Table 3 can be calculated, indicating that for every degree of temperature increase, the hydrolysis rate *k* will increase by a factor of 1.03 to 1.08 and half-lives will decrease accordingly. The impact of temperature decreases with increasing acyl-chain length. The presence of an oxo-side chain reduces the impact of temperature by approximately 0.01.

**Table 3:** Temperature-dependent factors for hydolysis rate *k* and half-life time *t* _*1/2*_ for a +1°C temperature increase derived from data by Yates *et al*. (2003).

| AHL | $Q_1$ for $k$ | $Q_1$ for $t_{1/2}$ |
| --- | --- | --- |
| C4 | 1.08 | 0.93 |
| C6 | 1.07 | 0.93 |
| C8 | 1.03 | 0.97 |
| C6-oxo | 1.06 | 0.94 |

To obtain the most representative data basis for further analysis, we combined the naturally relevant data obtained by Decho *et al*. [12] with the more chemically accurate, time-resolved data by Ziegler *et al*. [28] for C6 and C8. For both of these AHLs, the pH [12, 28] and temperature [22] influences on their abiotic decay have been established. Comparability of both datasets was ensured by accounting for the kinetic isotope effect of D_2_O compared to H_2_O and adjustment of the temperature by employing the coefficients from Table 3. Individual data points were then plotted and analysed as above, yielding linear regression equations with a very good fit (R^2^ > 0.95) as shown in Fig. 3 (detailed fit parameters specified in Table S6).

**Figure 3:**
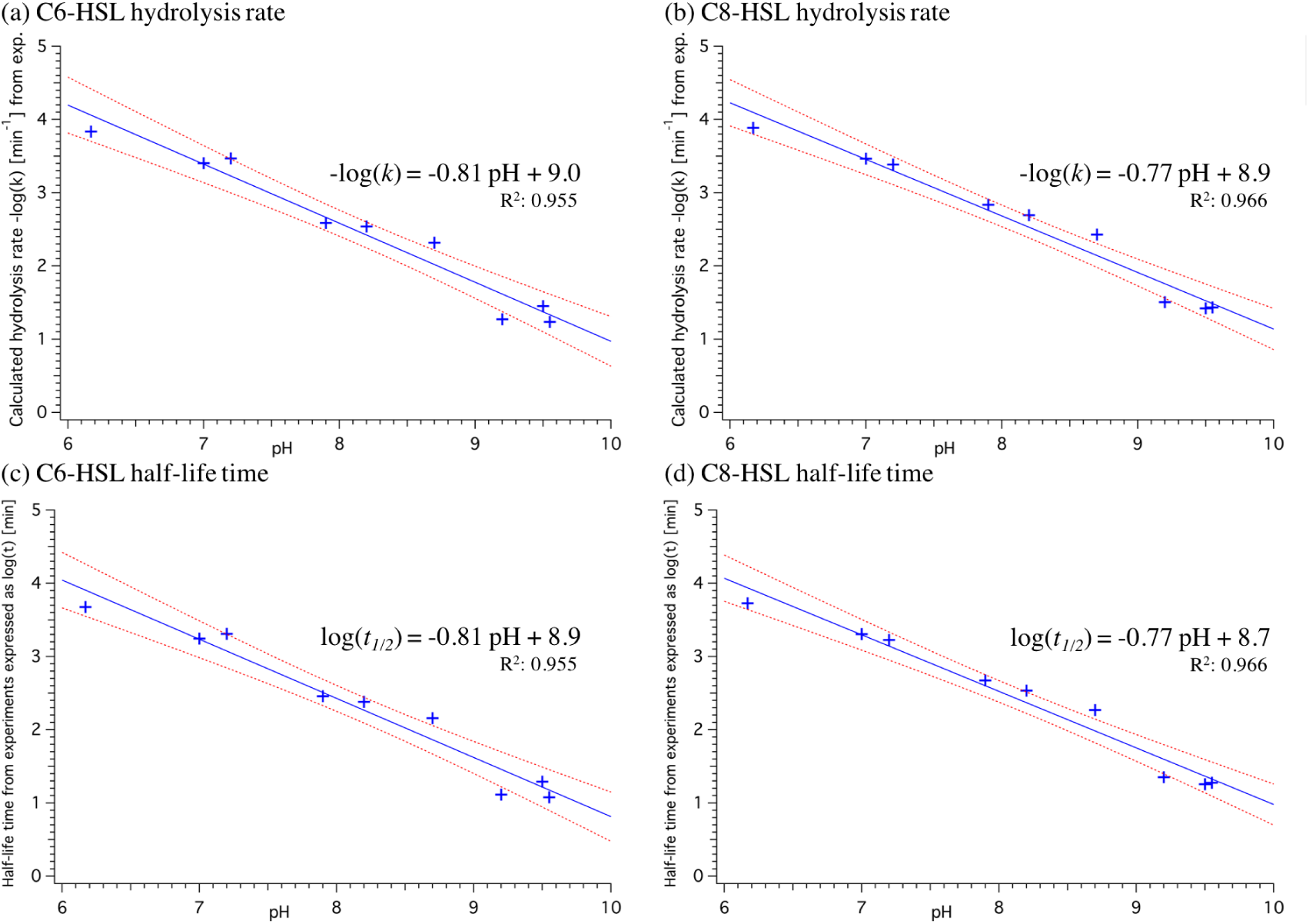
pH-dependence of hydrolysis rate *k* (top) and half-life time *t*_1*/*2_ (bottom) for C6- and C8-HSL with a pooled dataset including measurements from Decho *et al*. [12] and temperature adjusted data from Ziegler *et al*. [28] for which the kinetic isotope effect in D2O was accounted for. Linear least-square fit (blue line) yielded the respective equation. Red dashed lines indicate 95% confidence bands on fit and R2 value indicates goodness of fit (the closer to 1 the better). Standard deviation of fit coefficients is specified in Table S6.

### 3.2 AHL hydrolysis in current and future average conditions

In current average ocean sea-surface pH and temperature conditions, the hydrolysis rate *k* of C6- and C8-HSL is considerably faster by 0.70 × 10^−3^ and 0.75 × 10^−3^ per minute compared to future ocean conditions. This means that in average conditions predicted by the IPCC under a RCP8.5 ‘business-as-usual’ scenario for the year 2100, [1], the hydrolysis rate for these two AHLs will be 38% and 45% slower compared to today, respectively (Table 4). In turn, the half-life time of both AHLs will be increased in future by 61% for C6-HSL and 82% for C8-HSL compared to today. This equals an increase in half-life time by more than 4 or even more than 5 hours, respectively, compared to the half-life time in current conditions.

**Table 4:** Hydrolysis rate and half-life time of C6- and C8-HSL in average current and future conditions.

| AHL | Current average conditions:<br>16°C, pH 8.1 |  | Average conditions in the year 2100*:<br>20°C, pH 7.7 |  | Difference due to climate change |  | Relative change in future conditions compared to today |  |
| --- | --- | --- | --- | --- | --- | --- | --- | --- |
| | $k$<br>[10<sup>-3</sup> min<sup>-1</sup>] | $t_{1/2}$<br>[min] | $k$<br>[10<sup>-3</sup> min<sup>-1</sup>] | $t_{1/2}$<br>[min] | $\Delta k$<br>[10<sup>-3</sup> min<sup>-1</sup>] | $\Delta t_{1/2}$<br>[min] | $k$ | $t_{1/2}$ |
| C6 | 1.85 | 428 | 1.15 | 690 | -0.70 | 261 | -38% | +61% |
| C8 | 1.66 | 381 | 0.91 | 694 | -0.75 | 314 | -45% | +82% |

\* based on IPCC RCP8.5 business-as-usual scenario

The difference in hydrolysis rate/ half-life time between current and future average conditions also results in a noticeable difference in the decay of C6-HSL and C8-HSL over the course of 10 hours, as shown in Fig. 4. Due to climate change, there will be less abiotic hydrolysis of both AHLs. In average future conditions at pH 7.7 and 20°C, there will be 17.2% more C6-HSL and 21.0% more C8-HSL after 10 hours compared to current average ocean conditions. The concentration of C6 and C8-HSL reached in current conditions after 10 hours is only reached after more than 16 or 18 hours in future conditions, respectively, resulting in the chemical signals lasting for up to 8 hours longer.

**Figure 4:**
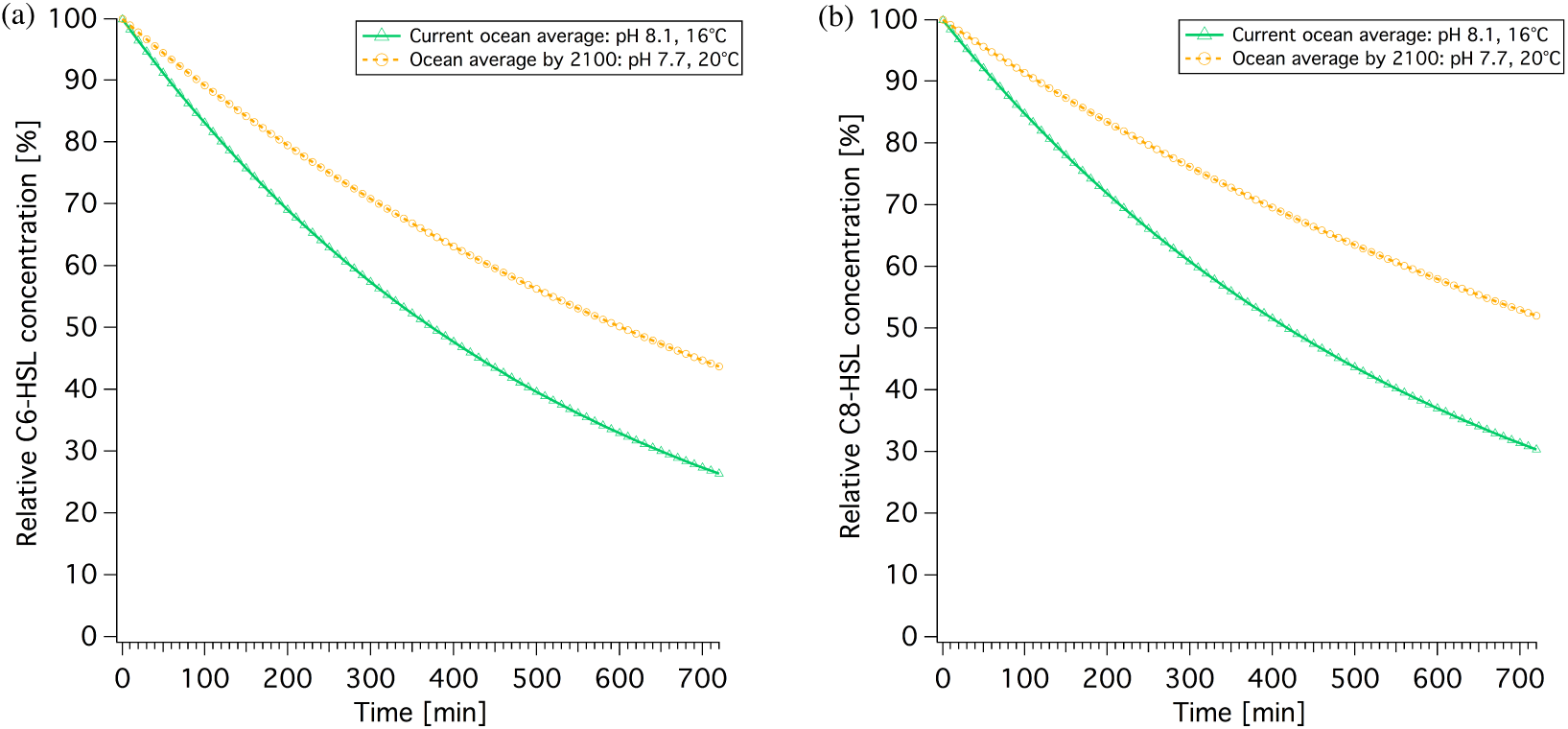
Current and future AHL concentrations over time for C6-HSL and C8-HSL. Decay is based on the respective hydrolysis rate stated in Table 4.

### 3.3 AHL hydrolysis dynamics in fluctuating conditions - quantification of natural variability

While average changes are important to gain an impression of the overall impact of ocean acidification and increased temperature, the natural variability within a system at different levels of spatial and temporal resolution can be equally important in order to obtain a holistic picture and understand baseline variability.

#### 3.3.1 Variability within the biofilm due to daily pH fluctuation

Measurements by Decho *et al*. [12] in marine stromatolite mats in the Bahamas revealed substantial daily pH fluctuations of up to 2.6 pH units despite an external stable pH of around 8. Assuming that a relative shift of the external pH (−0.4 units) would equally translate to the pH fluctuations within the biofilm allows a prediction of the impact of future pH-conditions (Fig. 5). Temperature was kept constant in this instance.

**Figure 5:**
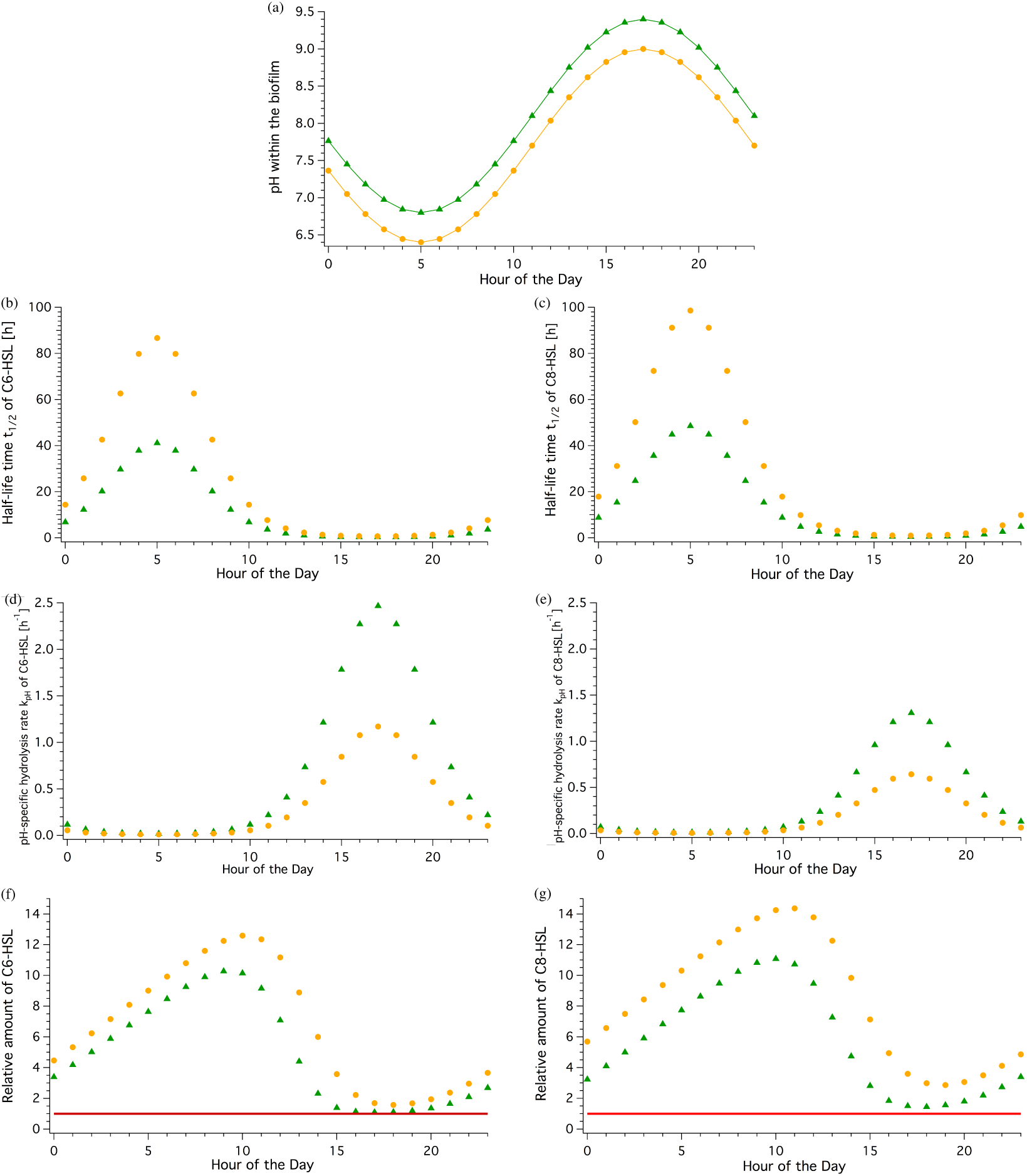
Current and future concentrations of C6-HSL and C8-HSL over a daily pH-cycle within a biofilm. (a) Periodically fluctuating pH-conditions based on Decho *et al*. [12] assuming stable temperature. (b) & (c) Half-life time in hours across a daily cycle for C6- and C8-HSL, respectively. (d) & (e) Hydrolysis rate *k*_*pH*_ over the course of the day for C6- and C8-HSL, respectively. (f) & (g) Relative amount of C6- and C8-HSL for every hour in a daily cycle. Amount produced is normalised to 1 (red line), so relative amount value reflects multitude of produced amount (accumulation).

Half-life time was greatest and hydrolysis rate slowest at 5:00 am in the morning, coinciding with the lowest pH value. Likewise, the lowest half-life time and fastest hydrolysis rate were observed at 17:00 in the afternoon when the highest pH is reached (green data points in Fig. 5).

Over the course of the day in current conditions, the half-life time of C6-HSL was found to range from over 41 hours in the early morning to as little as 19 minutes in the afternoon. For C8-HSL, *t*_1*/*2_ similarly ranged between 48.5 hours and 29 minutes. The hydrolysis rate displays the inverse trend ranging from 0.01 h^−1^ in the early morning to 2.47 h^−1^ in the afternoon for C6-HSL and a range from 0.01 h^−1^ to 1.31 h^−1^ for C8-HSL, respectively. Assuming a constant production (normalised to 1) and summing up produced and remaining AHL amounts taking the different hydrolysis rates into account, fluctuating daily AHL concentration patterns become apparent. In current conditions, the C6-HSL concentration reaches the highest level with 10.3 times the produced amount at 9:00 in the morning and drops to the lowest amount at 17:00 in the afternoon. For C8-HSL a similar pattern with slightly shifted timings (lag) is observed with a maximum exceeding 11 times the produced amount at 10:00 am and a minimum at 6:00 pm. This means that the AHLs accumulate to amounts over a magnitude higher than what is produced over the course of the night and into the morning before they degrade back to amounts close to the baseline level. While accumulation happens over a timeframe of 16 hours, degradation happens twice as quickly, within 8 hours.

In future conditions expected for the year 2100, half-life time and hydrolysis rate show the same patterns, coinciding with highest and lowest pH conditions as can be expected (Fig. 5, orange points). However, the linear shift of −0.4 pH units does not translate linearly, leading to more than double the half-life time at any given hour compared to the current conditions, and less than half the hydrolysis rate. This results in significantly higher levels of C6- and C8-HSL being present throughout under these future scenarios. C6-HSL accumulates for 16 hours to 12.6 times the amounts produced under current conditions, and is then degraded within 8 hours. Compared to current conditions, that’s 2.3 times the produced amount of C6-HSL at peak time in future conditions. Bacteria in future conditions could produce 18% less C6-HSL throughout the day to reach the same maximum concentration as in current conditions. For C8-HSL the differences for future compared to current conditions are even greater, with 14.4 times the produced amount at peak hour, 3.3 more than in current conditions. To achieve the same maximum peak concentration in future conditions, bacteria could produce 23% less C8-HSL throughout than in current conditions. Furthermore, the time at which maximum accumulation and lowest level of AHL is observed in future conditions is shifted by one hour for both AHLs.

The 16h accumulation and 8h degradation phases stay the same. Hence the reduction in pH due to ocean acidification can be expected to increase the baseline level of AHL concentration if the same level of production is maintained and shift the timing of the accumulation cycle.

#### 3.3.2 Seasonal variability based on the example of the Humber estuary conditions

Fluctuating conditions affecting habitats in coastal areas and estuaries, especially where there is significant tidal influence and/or fluvial input, were also found for the Humber estuary. The pH was found to vary between 7.2 and 8.4 without a clear seasonal pattern and mostly driven by tidal effects. Some very low pH values between pH 6 and 7 were measured early and late in the year, correlated to heavy rainfall events. Temperature, in contrast, had a clear seasonal trend, as expected, and could be fitted with a sinus equation with an average temperature of 10.99 (±0.07)°C and an amplitude of 6.5 (±0.1)°C (see Fig. 6 a & b). The pH and temperature adjusted half-life times and hydrolysis rates of C6-HSL calculated for each datapoint show the significant impact of the seasonal temperature pattern on these two parameters, but also reveal that there is a strong dependence on the pH causing large variability within a shorter than seasonal amount of time (days). Half-life time of C6-HSL throughout the year in the Humber estuary was found to be 23 hours on average, varying by ±13 hours due to seasonal influences (±57%). The hydrolysis rate was calculated to be on average 0.05 h^−1^, varying depending on season by ±0.024h^−1^ (±48%). Especially during the summer month the combined pH and high temperature conditions seem to cause fairly high hydrolysis rates (> 0.1h^−1^) compared to the rest of the year (Fig. 6d). When averaged across all data points for each month, the half-life time and hydrolysis rate showed significant differences across the year. Half-life time of C6-HSL in autumn and winter (Oct to Mar) exceeded 20 hours and was significantly longer than in spring or summer (April to September)(Fig. 6e, green bars). This was inversely reflected in the hydrolysis rate showing highest rates from April to September ranging between 0.05 and 0.08 h^−1^ (Fig. 6f, green bars). Shifting temperature by +4°C and pH by −0.4 units for every datapoint in line with IPCC predictions for conditions in 2100 results in significantly increased half-life times, which are on average 61% longer than those calculated for current conditions following the same seasonal pattern, and the hydrolysis rate in future conditions is on average 38% slower (Fig. 6e & f, orange bars).

**Figure 6:**
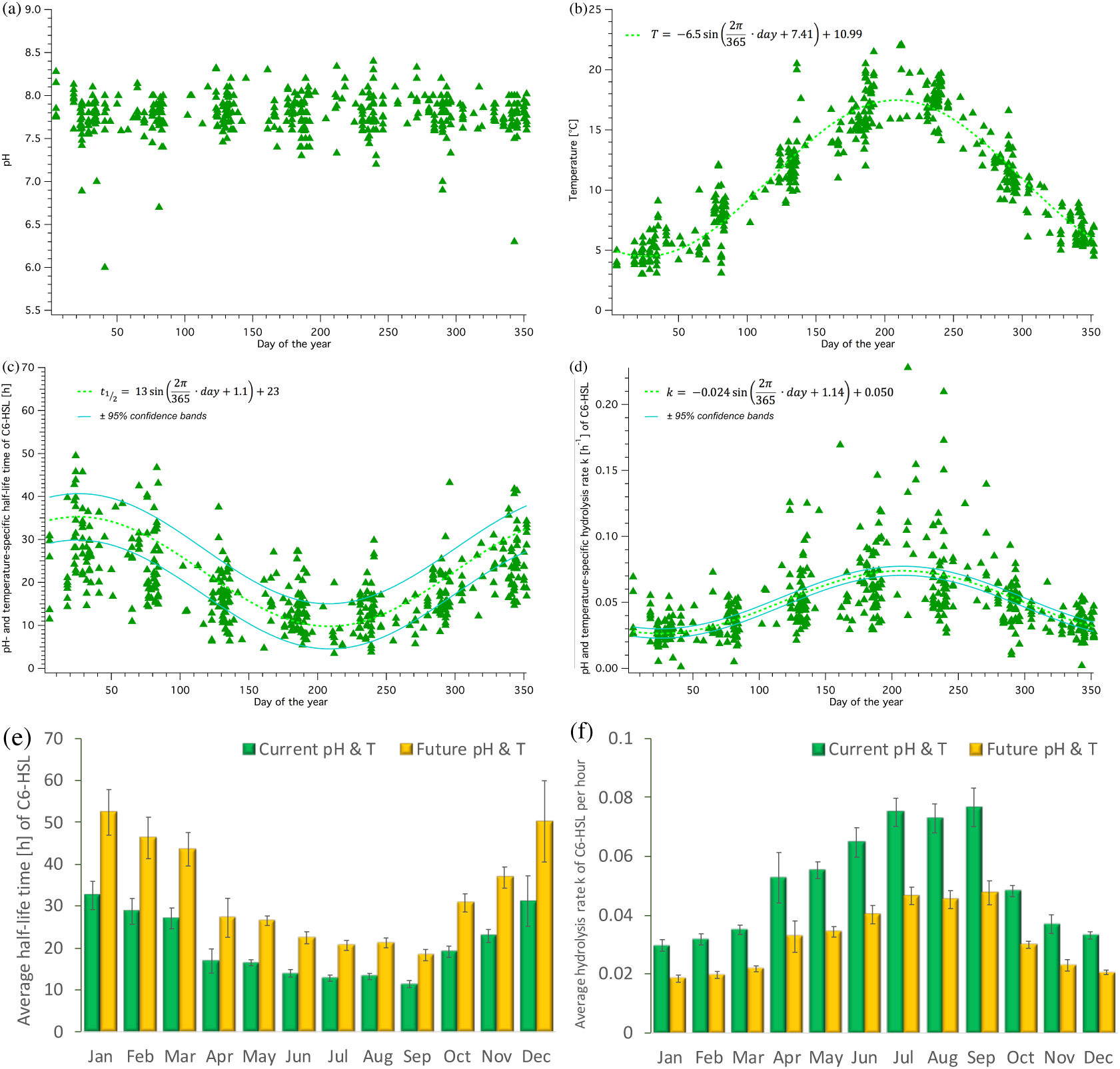
Seasonal fluctuations of pH, temperature, C6-HSL half-life time and hydrolysis rate over the year. (a) pH and (b) temperature measured within the Humber estuary (1995-2005) across the annual cycle. Trend of (c) half-life time and (d) hydrolysis rate of C6-HSL across the year assuming Humber conditions. Average (± SEM) monthly half-life time (e) and hydrolysis rate (f) for current (green) and future (yellow, based on average IPCC prediction of RCP8.5) Humber conditions.

#### 3.3.3 Combined seasonal and daily fluctuations with a perspective on future conditions

Seasonal differences in the water surrounding the biofilms with the AHL-producing bacteria are also potentially reflected inside the biofilm. To assess and visualise the impact of external pH and temperature conditions on the daily fluctuations within the biofilm for each month (including average, maximum and minimum conditions), the respective hydrolysis rates were calculated for C6-HSL based on equation (16) and the corresponding parameters determining *k*_*C*6_ from Fig. 3a as well as the respective temperature coefficient. Results are shown in Fig. 7. From January to April the impact of external factors was broadly comparable and highest pH and temperature conditions resulted in a hydrolysis rate of around 1 h^−1^ in the afternoon at peak pH within the biofilm. Minimum pH conditions at low and high temperatures resulted in very low hydrolysis rates. From May onwards the hydrolysis rates, especially in highest pH and temperature conditions, increase considerably, but there is also a larger variability of hydrolysis rates depending on the external conditions. Rates in November and December are lower again with less variability, similar to those in spring. It further becomes apparent that both, pH and temperature have a considerable impact on the hydrolysis rate within the biofilm.

**Figure 7:**
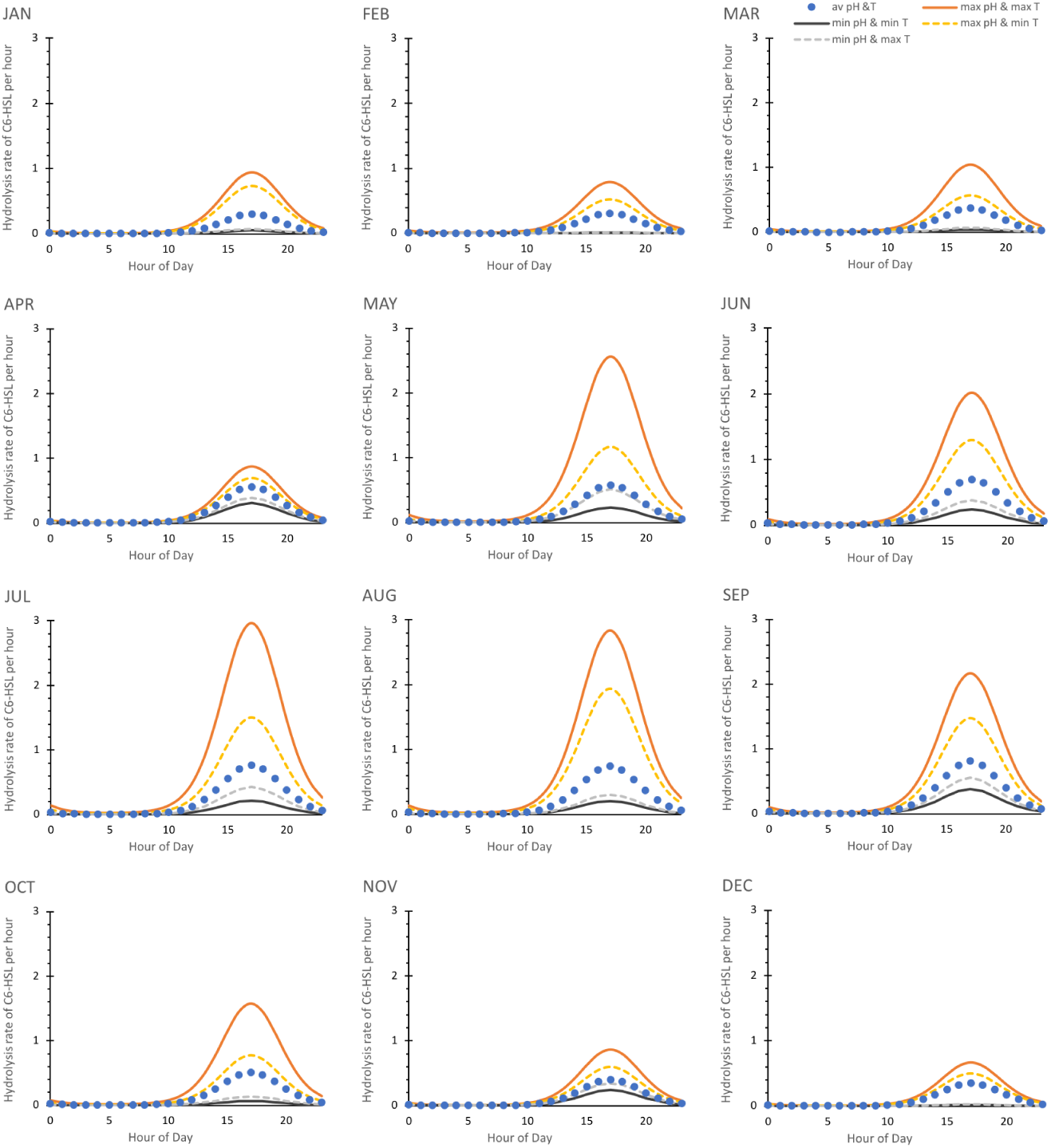
Daily fluctuation of C6-HSL hydrolysis rate for average, minimum and maximum pH and temperature conditions each month. Hydrolysis rate *k* is given in values per hour and calculated based on equation (16) and the seasonal average, maximum or minimum pH and temperature conditions of the Humber estuary dataset assuming that they translate unchanged to the biofilm as baseline conditions.

The seasonal effects on the daily dynamics of the C6-HSL hydrolysis rate within the biofilm will ultimately be reflected in the amount of C6-HSL that accumulates or degrades, as shown in Fig. 8. Relative levels of C6-HSL are highest in January and February, and lowest in July/August/September. During winter, C6-HSL amounts accumulating within the biofilm can exceed 20 times the amount of what is produced. In contrast, during summer peak C6-HSL only reaches levels of about 14 times the produced amount. This reflects a considerable seasonal variability of the AHL amount.

**Figure 8:**
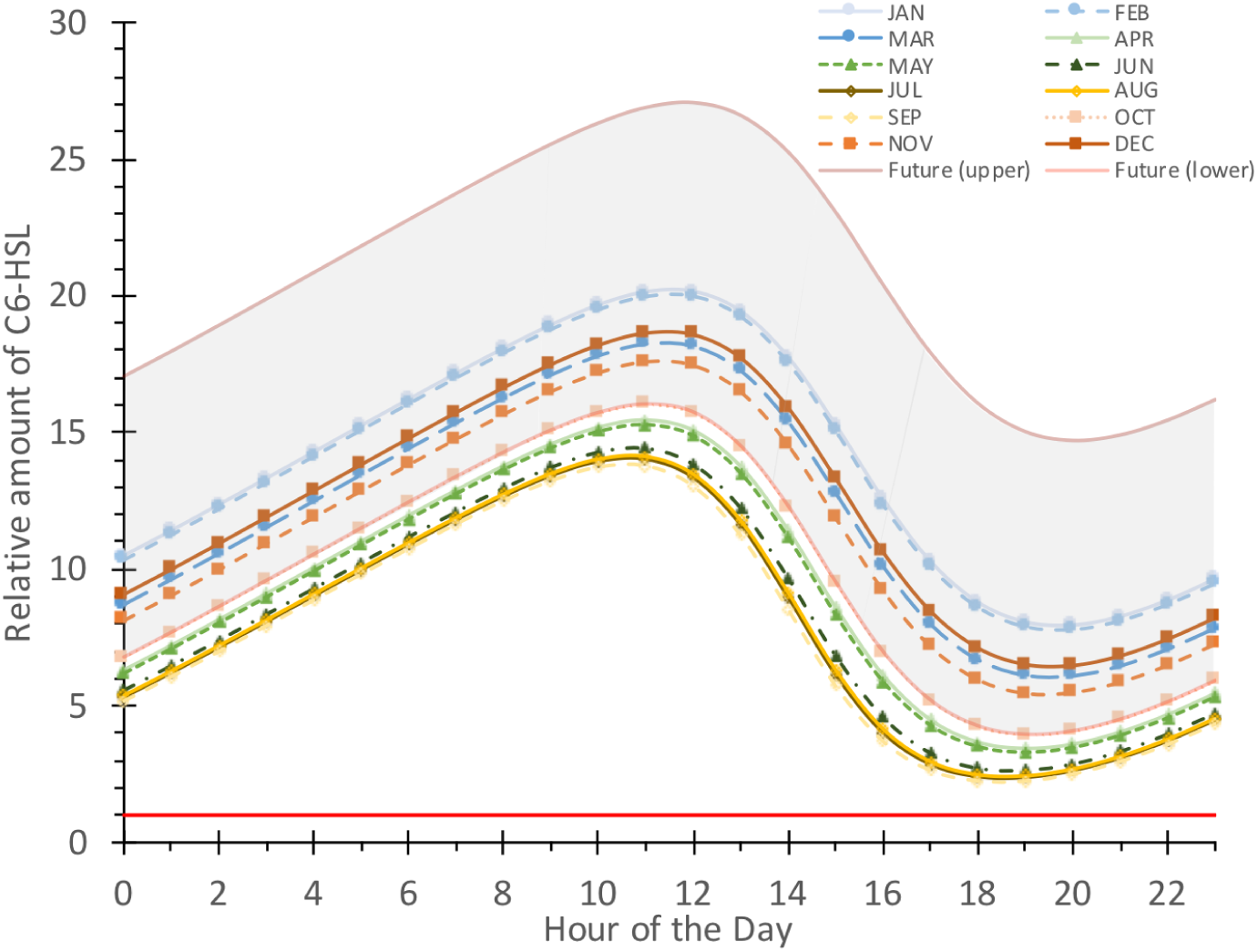
Comparison of daily fluctuating relative C6-HSL concentration for average pH and temperature conditions each month. Relative amount is calculated based on a normalised production of 1 (red line) and current conditions within the Humber estuary for every month (averaged). The projected future range of the fluctuation is shown in grey with the upper and lower boundaries representing January (upper) and September (lower) C6-HSL amounts calculated for conditions shifted by the IPCC RCP8.5 prediction (−0.4 pH, +4°C).

In addition to the variability in the accumulating and degraded amount, there is a considerable shift in timing when the maximum or minimum C6-HSL level is reached. In January and February, peak C6-HSL levels are reached at noon and minimal levels occur at 8pm. From March to December the maximum levels are already reached an hour earlier (11am) and degrade to the minimum within 9 hours in the case of March and December, or 8 hours to a minimum at 7pm in April to August, October and November. In September, the minimum level is reached already after 7 hours at 6pm. The differences in the timeframes of C6-HSL degradation highlights the considerable seasonal impact on the dynamics of this signalling system.

Placing this seasonal range in the context of future conditions by adjusting the relevant pH and temperature values relative to the IPCC RCP8.5 prediction (−0.4 pH, +4°C) yields a substantial shift of the C6-HSL amounts, which are found to accumulate at even higher levels, and up to 27 times the levels produced amount during winter and 16 times the produced amount in summer, the latter being comparable to October levels under current conditions. Minimum levels are also raised compared to current conditions. Timings were found to be affected by seasonal differences, as observed for the current conditions.

These results highlight the substantial impact of climate change on the dynamics of AHLs like C6-HSL which far exceed naturally occurring variation found in current conditions.

## 4 DISCUSSION

The key purpose of this study was to investigate theoretically how the degradation of AHLs is affected by abiotic environmental changes. We established a numerical relationship between pH and AHL hydrolysis rate/ half-life time and calculated temperature coefficients for all relevant conditions based on collated published data. By comparing the impact of pH and temperature on AHL concentration individually and combined at different timescales, this study reveals that natural daily and seasonal, as well as projected climate change associated abiotic changes, all have the potential to considerably influence the dynamics of AHLs in biofilms and thus impact biofilm form and function.

### The daily rhythm of AHL dynamics driven by pH and the importance of other influencing factors

Within a daily timeframe, a cycle of accumulation and degradation of AHLs occurs in a rhythmic pattern arising from the impact of the natural pH fluctuations inside the biofilm, based on the hydrolysis rate. Higher pH in the afternoon, thought to be caused by the photosynthetic activity of biofilm-associated phototrophic organisms, [12] leads to a sharp increase in the hydrolysis rate and in turn to an up to 3 times faster degradation of the AHL molecules (Fig. 5 d&e, Fig. 7). This pH-driven dynamics can be enhanced or reduced to a small extent by temperature (Fig. 7). Declining pH during hours with little or no light, and hence little photosynthetic activity within the biofilm, slows down the hydrolysis rates and results in much longer half-live times of the AHL molecules during the early hours prior to sunrise (Fig. 5 b&c). AHL concentrations reach their peak in the late morning and their lowest level in the late afternoon (Fig. 5 f&g, Fig. 8). This dynamic does not directly mirror the timing of lowest and highest pH, but the highly increased hydrolysis rate at peak pH diminishes the available AHLs very quickly so that the lowest AHL concentrations occur shortly after the pH maximum.

To validate our theoretical results, we compared the difference in concentration of C8-AHL between 6am and 5pm, the times when actual measurements were taken by Decho and co-workers [12]. They analytically determined the C8-AHL concentration in the morning to be 6.5 ± 0.8 ppb, 1.8 times (± 0.5) higher than in the afternoon (3.6 ± 0.9 ppb) [12]. Our calculations yield a 5.3-times higher concentration in the morning than in the evening for the same times, exceeding the experimental results approximately 2-fold. There are a number of potential reasons for this theoretical overestimation of the AHL difference in our approach: inconsistent AHL production, deviations of the dynamics-determining parameters, and additional AHL degradation through other mechanisms. Firstly, for our calculation we assumed a consistent level of AHL production throughout the day. It is, however, likely that AHL production and excretion by bacteria occur inconsistently and locally [26] as it forms part of their cell-cell communication. This may vary across a daily cycle and is known to potentially depend on other internal and external factors, such as cell density and distribution, composition and physical characteristics of the surrounding medium, available carbon sources and oxygen limitation amongst many others [33]. Infrequent production would cause a reduced amount of accumulating AHLs and therefore a smaller difference. Secondly, less pronounced dynamics of AHLs in nature could also be caused through deviations from the parameters used in our calculations, particularly the hydrolysis rate. An underestimation of the naturally occurring hydrolysis rate could have caused the greater amounts of AHLs accumulating in our calculations. The hydrolysis rate we employ is based on an averaged value derived from abiotic laboratory experiments published by Decho *et al*. [12] and Ziegler *et al*. [28]. This rate does not take into account any naturally occurring temperature fluctuations, which might have influenced the natural hydrolysis and hence analytically measured AHL concentrations. Unlike for pH, temperature fluctuations were not quantified and reported across the daily timeframe for the experimental data, limiting our possibility to take them into account in this calculation used for comparison. The laboratory experiments underlying the hydrolysis rate were further conducted with different chemical methods (extraction + GC/MS vs. NMR), which is why we combined and averaged the respective results for a more representative rate. It might be that the slightly faster rates determined using GC/MS [12] are closer to those in the field compared to the rates obtained through NMR measurements in D_2_O and subsequent conversion [28] (see Table 1 for comparison). It is, however, interesting to note that other experiments aiming to identify degradation rates of AHLs in seawater found significantly lower rates [19, 23]. Tait & Havenhand determined the degradation of C6-HSL and C8-HSL in seawater at 18°C to be 1.5 (±0.2)% /hour and 1.0 (±0.5)% /hour, respectively. [19] These rates are by a factor of 6 to 7 slower than those reported by Decho and coworkers for pH 8.2, but similar to those reported by Hmelo & van Mooy (pH 7.9), who suggest that AHL degradation in seawater might be slower than in non-marine media [23]. An underestimation of the hydrolysis rate in our approach is therefore unlikely to explain the discrepancies between our calculated difference and the naturally observed difference. Thirdly, the detectable amount of AHL and its accumulation could be affected by other diminishing factors such as leaking of AHL from the biofilm, enzymatic AHL degradation or metabolisation by other organisms [10]. This additional loss could account for the 66% smaller AHL difference observed in nature compared to our calculated result. While leaking can be expected to be comparably low due to the protective EPS matrix of natural biofilms [33], it does occur to some extent [17] as evidenced by several AHL-induced interactions with settling macro-organisms [10, 19] (for overview). These interactions require sensing of AHLs by the macro-organism in the water column from a distance to the biofilm. Also, AHLs with shorter chain lengths (e.g. C4, C6) are less hydrophobic and consequently diffuse more rapidly into the surrounding than longer-chain AHLs. [17, 33] Besides leakage, biotic degradation of AHLs through enzymes is a factor that could explain a substantial part of the difference between our calculations and the AHL levels as it is known to play a key role in bacterial biofilms. [33] Hmelo & van Mooy found 54% of C6-HSL degradation in seawater to be likely caused by enzymatic activity [23]. Because AHLs serve as fundamental cell-cell communication in many bacteria, disruption of this communication pathway by quenching AHLs enzymatically provides competitive benefits for other bacteria and is in fact widespread [33, 34]. Two major types of AHL quenching enzymes have been described, lactonases and acylases, [35] which hydrolyse the lactone ring [36, 37] or cleave the amide bond of the AHL ‘tail’ [38], respectively. While these enzymes work across a wide range of environmental conditions, [39] some were found to follow a steep pH-dependent optimum curve [40], potentially adding to the complexity of the pH-dependent daily dynamics of AHLs.

Despite the likely influence of other AHL degrading factors in nature as shown by the direct comparison, our investigation reveals that abiotic AHL degradation through hydrolysis linked to a daily cyclic pH pattern plays an important role, yielding results in the same order of magnitude as comparable experimental measurements. Our results overestimate the difference by a factor that matches the 50 to 60% AHL observed to be lost through enzymatic degradation [23]. In turn, this means that abiotic hydrolysis accounts for at least 1/3 or more of the observable dynamics. For subsequent interpretation of our results, the importance of other influencing factors, however, has always to be taken into consideration. It also has to be noted that there is an apparent lack of biofilm parameters and abiotic conditions that are monitored continuously or regularly for a daily timeframe and local context, e.g. near sediment or rocky colonised surfaces. We therefore suggest a focus of future measurements on the daily patterns of natural pH and temperature in direct relation to AHL concentrations with hourly intervals within the natural habitat of interest, for example surface biofilms on sediment or rocky substrate.

### Seasonal impacts on AHL dynamics driven by temperature

AHL hydrolysis rate and half-life time showed a clear seasonal pattern across the year with results in hydrolysis rate varying by 48% and half-life time by 57% largely due to the temperature influence. Significantly higher hydrolysis rates in spring and summer, and, in contrast, half-life time exceeding 20 hours in autumn and winter, clearly mimic the temperature pattern. The change in hydrolysis rate between winter and summer exceeds a factor of 2, suggesting that seasonal conditions impact AHL dynamics in a way that is likely reflected in the overall dynamics, despite other influences. Combining seasonal and daily fluctuations in pH and temperature revealed that seasonal differences are reflected in the daily patterns and subsequently cause a shift in the daily cycle. In summer, AHL levels accumulate to only 70% of winter levels, taking an hour longer to do so and becoming degraded within only 7 hours, so one hour quicker than in winter. In addition, maximum AHL concentration in summer is reached two hours earlier in the day than in winter, shifting the timing of the cycle.

Our calculations for combined seasonal and daily dynamics assume a direct translation and addition of external conditions to the internal conditions within the biofilm. This means that external temperature was assumed to represent biofilm temperature and the external pH at any given date was used as the midline point for the biofilm-internal pH curve modelled with an amplitude of 1.3 across the day based on Decho *et al*. [12]. External pH and temperature changes might, however, be compensated by the biofilm-surrounding chemical matrix of EPS, which is assumed to buffer pH fluctuations [31] and extreme temperatures [41]. To what extent biofilms are able to actually compensate external abiotic conditions, however, is currently poorly understood and requires dedicated experiments, which simultaneously and systematically measure external and internal pH and temperature gradients in situ. Due to the afore mentioned other influencing factors, small seasonal differences indicated in our calculations need to be interpreted with caution. In a recent field study assessing the concentration of C8, C10 and C12-HSL in surface sediment of an intertidal mudflat, Stock *et al*. found AHL concentrations in samples from February and April to not differ significantly. [14] This contrasts the small but significant theoretical increase in average hydrolysis rate we obtained for April compared to February based on the data for the Humber estuary assuming similar seasonal patterns of both estuaries. Similar AHL levels for February and April further fits with the very similar daily hydrolysis rate profiles we obtained. To actually assess season-dependent AHL dynamics, further sampling over the summer, autumn and winter months would be required. The substantial differences between summer and winter we obtained, however, do suggest the potential for significant dynamics differences between these two seasons.

### pH and temperature as combined factors - enhancing or compensating effects depend on the timeframe

While pH changes dominate AHL dynamics within a daily timeframe, we observed temperature to particularly influence AHL degradation patterns in a seasonal context. Depending of the combination of these two factors, however, the hydrolysis rate can be sped up or slowed down. An increase in temperature increases the hydrolysis rate [22]. Higher pH also leads to a higher hydrolysis rate [12, 28]. Highest hydrolysis rates and consequently fastest degradation of AHLs can therefore be expected in the late afternoon and early evening during the summer months. In contrast, lowest hydrolysis rates and almost no degradation can be assumed for night and early hours in winter. These patterns can be observed as expected from our calculations of daily hydrolysis rates for each month (Fig. 7). The combined effect of temperature and pH is therefore clearly time-dependent on a daily and seasonal scale due to the corresponding natural fluctuations.

Climate change is predicted to result in higher temperatures and lower pH conditions [1]. The temperature-associated increase in hydrolysis rate is opposed by a pH-related reduction, which might result in effects cancelling each other out. However, our results reveal that the effect of pH exceeds the effect of temperature, resulting in a clear reduction in AHL hydrolysis and hence increased AHL concentrations for any of the future scenarios calculated.

### Climate change impacts - small average changes in the context of large natural abiotic fluctuations do matter for AHL dynamics

Looking at the impact of predicted average climate change related reduction in ocean pH and increase in sea surface temperature revealed an overall decrease in the hydrolysis rate of C6- and C8-AHLs in future oceans. This results in higher levels of the AHLs being present for longer in the environment (Fig. 4). Combining daily, seasonal and future parameters also clearly indicates the impact on AHL dynamics across these different timescales (Fig. 8). Future average changes in temperature and pH might seem small compared to the natural range of these parameters (+4°C compared to a natural seasonal temperature range of 13°C (31%), −0.4 pH compared to a daily pH range of 2.6 (15%)). But, while reflecting the daily and seasonal patterns, the future scenario results in even higher levels of C6-HSL, reaching more than 1.4 times the levels present under current conditions, and causes levels to never fall below current October levels by exceeding current winter levels by more than 30%.

The buffering of external conditions by the biofilm discussed previously and potential limitations due to our assumption of a direct translation of external factors to biofilm-internal conditions also apply in the context of future conditions. We further applied the projected average future changes in pH and temperature directly to the current natural ranges, resulting in a shifted range. An increasing number of studies, however, indicates that pH conditions are not only expected to shift but also considerably increase in variability [42], emphasising pH extremes. In addition, marine heatwaves are predicted to become more frequent and last longer. [1] Our results might therefore simplify and potentially underestimate the influence of future ocean conditions.

### Applicability of results to other AHLs

We focussed in this study on C6 and C8-HSL due to their documented presence and functions in marine biofilms [8, 12, 14] and the availability of sufficient data to determine pH and temperature impacts numerically. It is, however, important to note that the hydrolysis rate and the extent of pH and temperature influence depend on the chain-length of the AHL, [12, 22, 28] which is also reflected in our results (Tables 1 and 2). Shorter chain AHLs, namely C4 and C6-HSL, degrade faster with higher hydrolysis rates than AHLs with side chains of 8 or more carbons. [12, 22, 33] AHLs with a 3-oxo substitution, in contrast, degrade faster than their unsubstituted counterparts [22, 23] due to an additional abiotic degradation pathway via a Claisen-like condensation to tetramic acids [43]. For long-chain AHLs with longer half-life times, it can therefore be assumed that abiotic hydrolysis plays a minor role in signal termination and that most of these signals are degraded through enzymes to ensure termination of the signal within a relevant timeframe. The impact of fluctuating conditions and the resulting daily and seasonal dynamics shown here for C6 and C8-HSL may therefore not be as pronounced for longer-chain AHLs. But impacts of pH and temperature might be indirectly reflected in AHL concentrations as they might influence the kinetics of degrading enzymes. [40] AHL-quenching enzymes AiiA & Est isolated from a *Altererythrobacter sp*. strain from a marine beach (Red Sea) were found to actively cleave 3-oxo-C12-HSL in pH conditions between pH 5-10 or pH7 to >10, respectively. [40] However, optimum quenching activity was reached at pH 8 or 9 with significant reductions in activity for pH *<* 8. [40] A reduction by on average 0.4 pH units with ocean acidification could result in approx. 25% reduction in enzyme activity based on extrapolation of the data by Wang *et al*. [40]. This adds another layer of complexity by potentially enhancing the observed higher AHL concentrations in future conditions due to reduced quenching. Interestingly, our results also reveal that the extent of the impact of future conditions on abiotic hydrolysis rate and half-life time and consequently on the AHL concentration increases with increasing chain length. This is likely caused by the reduced compensating impact of the temperature influence on the hydrolysis rate (less acceleration) in relation to the impact of pH (reduces *k*), as longer AHLs are also less sensitive to elevated temperatures. [22, 33]. This results in a greater increase in concentration of AHLs with longer chains compared to shorter chain ones subjected to the same pH change. To more conclusively understand and estimate the impact of future conditions as well as natural fluctuations on AHL dynamics across the range of chain lengths and un-/substituted molecules, measurements of abiotic and biotic degradation under set environmental parameters need to be conducted.

### Biological and wider implications of AHL dynamics in current and future oceans

In the context of the substantial current fluctuations in AHL concentrations on daily and seasonal timescales, the impact of future ocean conditions shown in our results poses the question how an overall increase in concentrations and a change in timing of the AHL peak may affect marine, coastal and estuarine biofilms and their functioning.

For bacteria-bacteria interactions, the AHL communication system is finely tuned with AHL threshold concentrations for bacterial growth and adhesion ranging from 10 ng/L to 10 *µ*g/L (0.5-0.3 pM to nM) depending on biofilm composition and bacteria [44]. If higher AHL concentrations will prevail for longer in future conditions, as suggested by our results, bacteria would benefit, because less of the respective AHL needs to be produced to achieve the same threshold within the same timeframe. Likewise, if production remains unchanged, threshold concentrations would be reached faster or with a lower cell density, and the signal would be able to travel for a longer distance from the source, [23] making AHL-signalling more efficient. These potential impacts of future conditions were also hypothesised by Hmelo. [25] The enhanced longevity of AHL signals might boost biofilm formation, biofilm growth through enhanced bacterial cell growth and replication, and bacterial EPS and enzyme production [8, 25]. The range of the daily dynamics of C6 and C8-AHL in our study exceeds a factor of 10, which is even further enhanced in future conditions. Assuming that bacteria operate close to their concentration thresholds to maintain meaningful signalling, the daily cycle could lead to times during which AHL-signalling is facilitated (night and early morning) or prevented (afternoon) due to the conditions within the biofilm caused by the autotrophic co-habiting organisms. A similar conclusion on the possibility of AHL being involved in the timing of interactions was also reached by Hmelo [25] and Decho *et al*. [12, 45] with the latter establishing natural concentration differences across the daily cycle close to threshold concentrations in a range of 13 pmol/ g dry sediment for C8-HSL and 3.8 nmol/ g dry sediment for C10-HSL [12]. However, the shift to the cycle’s timing by an hour due to future conditions, as identified in our study, would likely have very limited impact, given that the current natural seasonal changes affect peak AHL times by up to two hours as discussed above.

Apart from enhancing the bacteria-bacteria interactions, higher and more stable AHL concentrations would also impact other interactions of importance in a biofilm context. Greater signalling power of C10-AHL, for example, could boost the formation of diatom-biofilms, as it has been shown to promote chlorophyll a concentrations and diatom-derived EPS production [16]. But threshold concentrations required to trigger increased carbohydrate levels in diatom biofilms (0.1 mg/L; 0.4 *µ*M) or enhanced diatom growth (1 mg/L; 4 *µ*M) are an order of magnitude higher than concentrations triggering bacteria-bacteria interactions. [16, 44] With our results suggesting a maximum increase of AHL concentration by approx. 20%, it is likely that these changes due to future ocean conditions alone will not impact these interactions substantially. However, they might enhance AHL accumulation at key times within the daily and seasonal cycles and thereby act synergistically to considerably strengthen and/ or prolong the signal. The same applies for interactions with macro-organisms. Concentrations of around 5 *µ*M necessary to induce the settlement of cypris larvae of *Balanus improvisus* [19] and more than 100 *µ*M to trigger exploratory behaviour of the polychaete *H. elegans* [20] might be exceeded earlier, at a lower bacteria density or reach further into the water column and hence trigger more settlement of macro-organisms due to signal enhancement through future conditions.

Future prolongation of signal life-span might, however, also poses issues: the short chain AHLs used as a form of short-messaging system in bacterial biofilms [25, 33] would not degrade as readily under future conditions and hence become less suitable for instant messaging. This might also affect the ratio of short- and long-chain AHLs in mixtures, which is hypothesised to play a role in complex settlement interactions with zoospores. [13] Due to their important role in the establishment of biofouling communities, higher AHL concentrations sustained for longer might also make biofouling of surfaces more common. [8, 25]

Signalling via AHLs is involved in fundamental biogeochemical and ecological processes in marine ecosystems, such as the remineralisation, dissolution or disaggregation of sinking particulate organic carbon, nutrient cycling, initial colonisation of surfaces and settlement of marine organisms (see Hmelo [25] for an overview). Changes to these processes would be of global significance. And we hypothesise that there is another fundamental ecosystem service likely to be affected by changes to AHL dynamics: biologically-mediated sediment stabilisation. Marine biofilms, in particular the EPS they produce, and the presence of vegetation and/or bioturbating organisms have been established as key factors in sediment stabilisation within coastal and especially tidal marine ecosystems and estuaries, such as saltmarsh and mudflat habitats. [46] AHLs are known to be of great importance in mediating biofilm communities, for example by inducing growth of diatoms [16] or boosting EPS production [47], and mediate the interactions with associated organisms like macroalgae [17] and bioturbating worms [20]. We therefore suggest that AHLs could be crucial mediators and quantitative changes to AHL concentrations can affect the sediment stability and thus highlight the need for more work to fully explore these important impacts.

## 5 Conclusion

Our study reveals that pH- and temperature-dependent abiotic hydrolysis of the key bacterial chemical signal class of AHLs leads to substantial theoretical dynamics of these important chemical signals in biofilms across daily and seasonal timescales. The work additionally highlights how these variations are amplified by a switch to projected future conditions caused by climate change. Our results indicate the importance of these abiotic drivers in the context of current natural fluctuations and other biotic influences on the AHL dynamics, showing that future ocean conditions likely result in higher AHL concentrations being present for longer, but within similar daily and seasonal cycles. The chemical dynamics of AHLs on different timescales could lead to changes in the timing of AHL-mediated processes and associated behaviours like the settlement of micro- and macro-fouling organisms. Future changes might not only enhance settlement, but also increase sediment stability by impacting estuarine biofilms. However, more detailed studies on the buffering capacity of biofilms with regards to external conditions on daily and seasonal timescales need to be conducted. The natural dynamics and importance of enzymatic degradation in relation to abiotic hydrolytic degradation in intertidal and estuarine biofilms need to be established for the full range of relevant AHLs with different chain lengths that are present in those biofilms. Direct links between AHLs and sediment stability due to cohesion through biofilms remain to be established.

## 6 Acknowledgements

This work was funded by ERC-2016-COG GEOSTICK (Project ID: 725955) and CCR acknowledges funding through a University of Hull Vice-Chancellor Research Fellowship.

## 7 SUPPLEMENTARY

**Figure S1:**
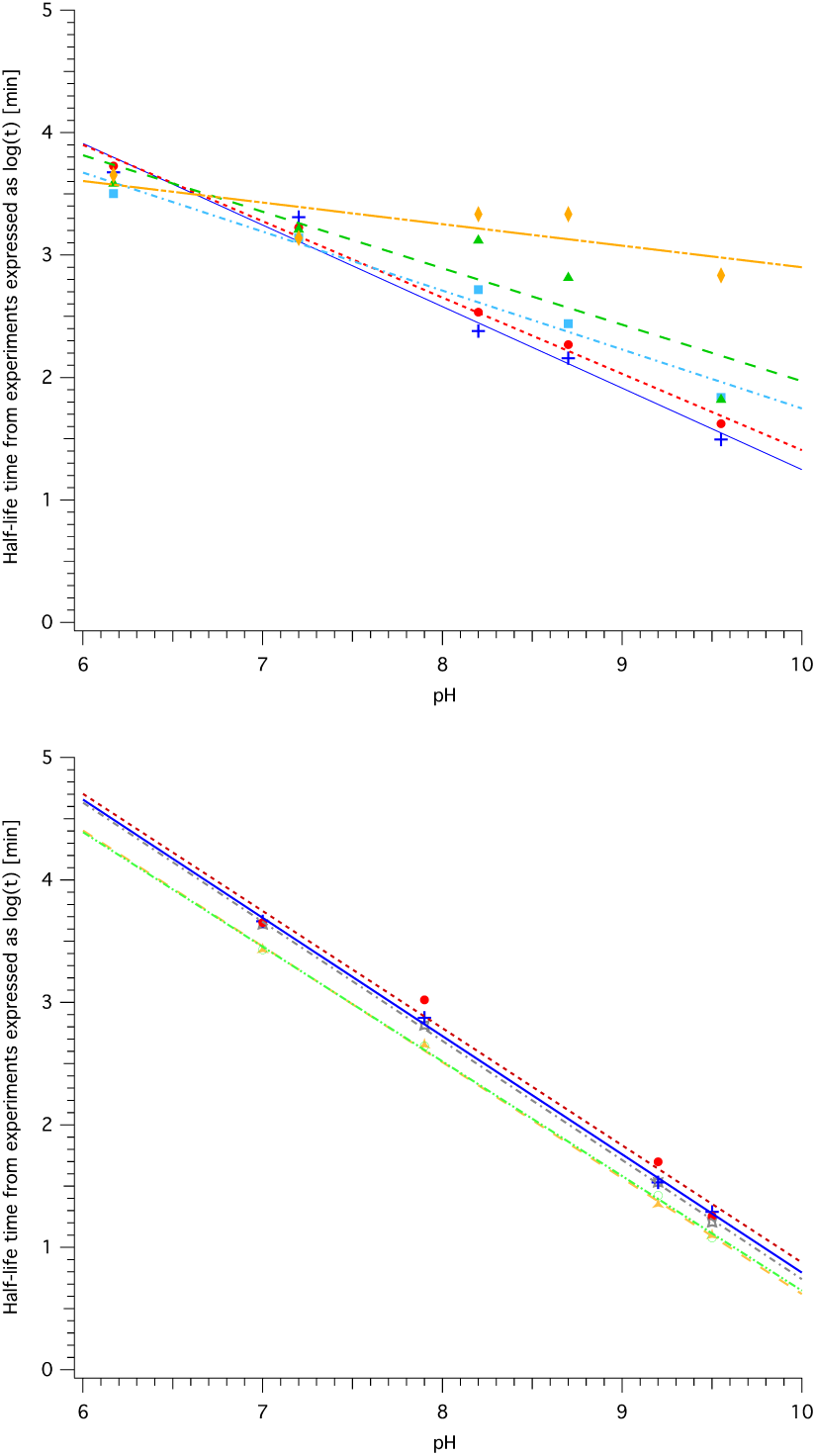
pH-dependence of half-life time *t*_1*/*2_ for different AHLs. Based on data published by Decho *et al*. [12] (left) for C6 (blue, cross), C8 (red, circles), C10 (light blue, squares), C12 (green, triangles) and C14 (orange, diamonds) and data published by Ziegler *et al*. [28] (right) for C4 (grey, star), C6 (blue, cross), C8 (red, circles), C6-oxo (orange, triangle) and C8-oxo (light green, open circles). Coefficients for linear least-square fit following the equation *log*(*t*) = *b × pH* + *D* are detailed in Table 2.

**Figure S2:**
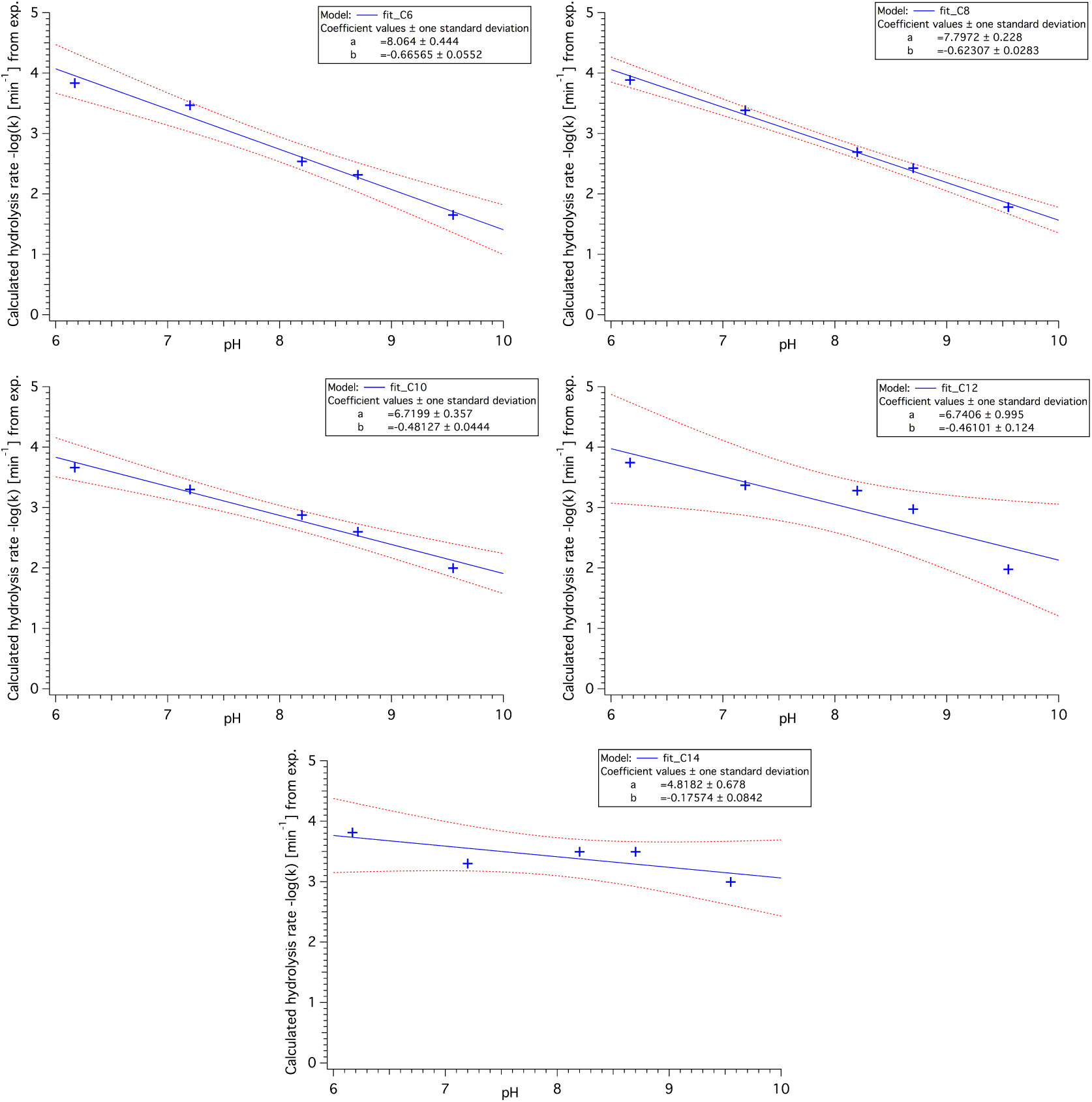
Hydrolysis rate kpH – pH - based on experimental values at 26°C by Decho.

**Figure S3:**
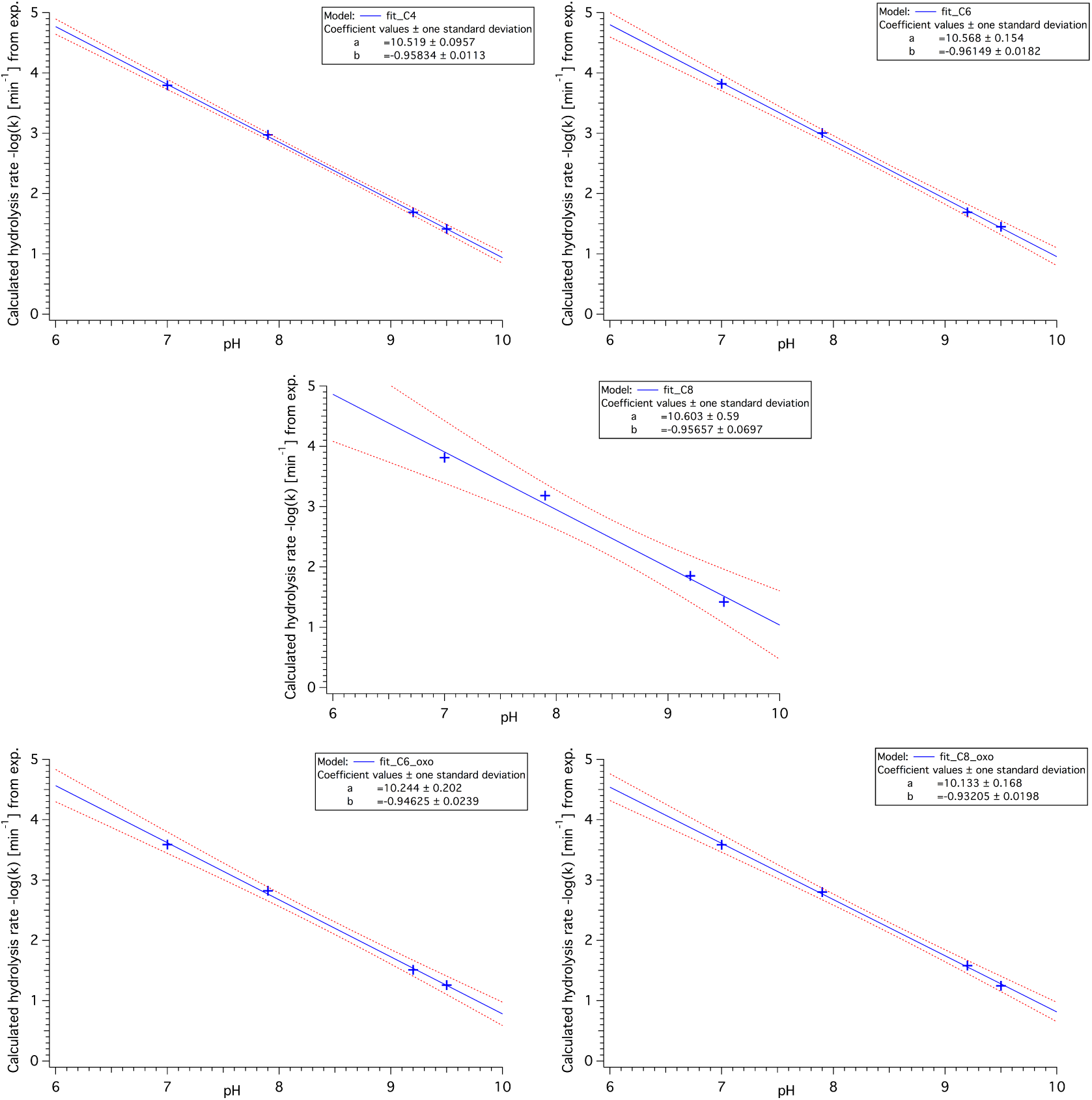
Hydrolysis rate kpH – pH - based on experimental values at 22°C by Ziegler.

**Figure S4:**
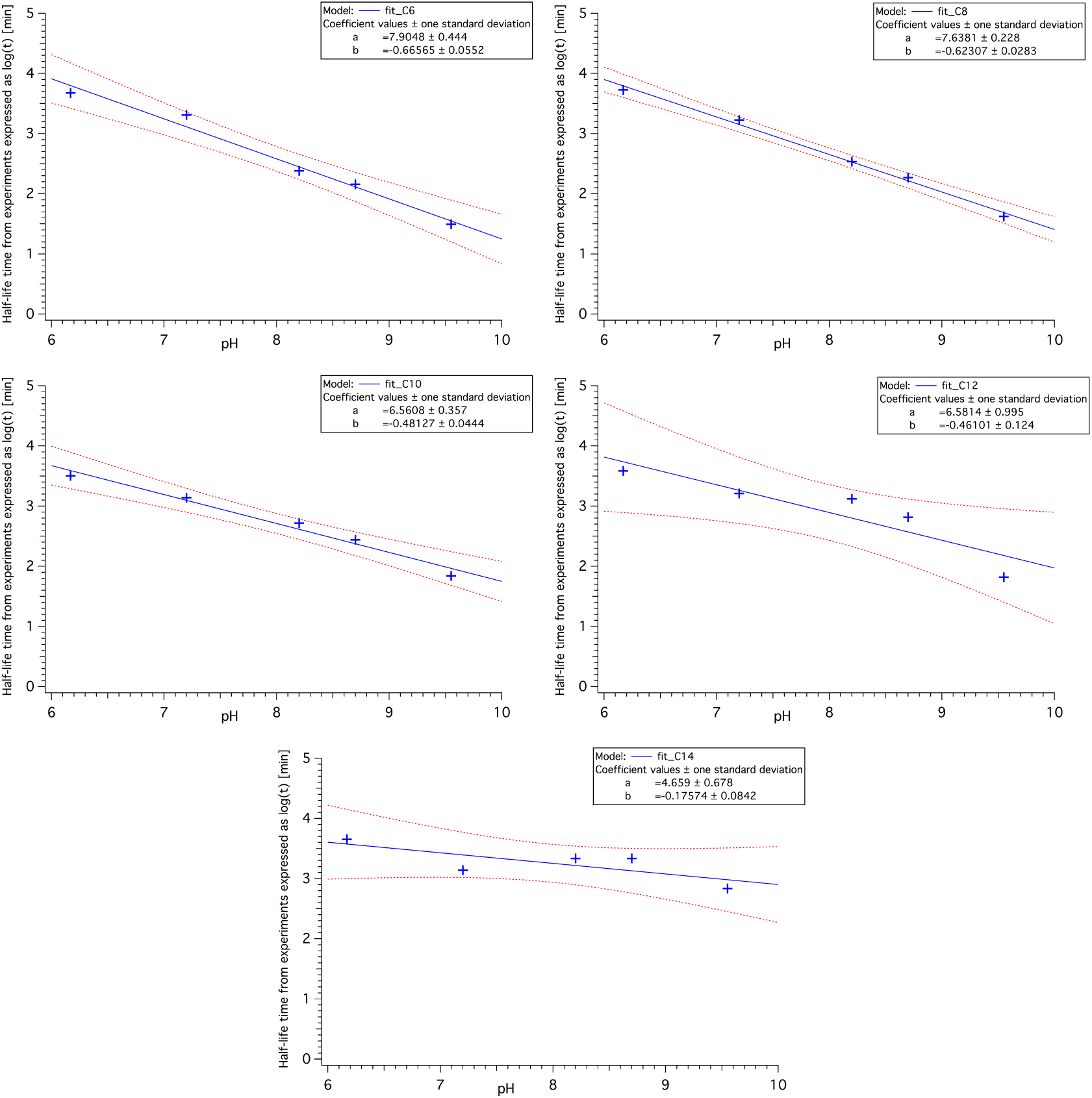
Half-life time – pH - based on experimental values at 26°C by Decho.

**Figure S5:**
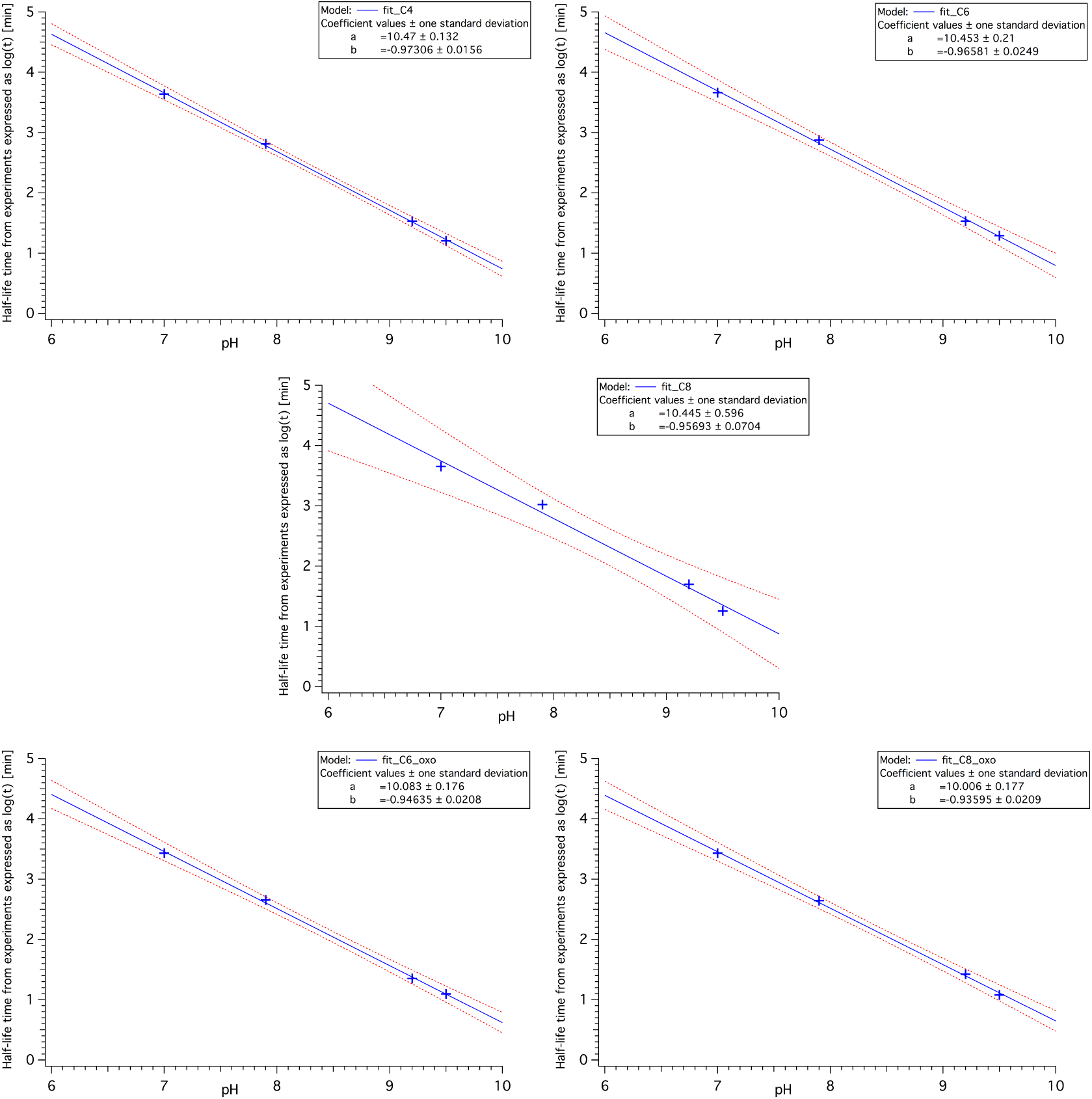
Half-life time – pH - based on experimental values at 22°C by Ziegler.

**Table S6:** Coefficients (± SD) of the relationship between hydrolysis rate *k* or half-life time *t* _*1/2*_ and pH for the combined dataset adjusted to H_2_O and 26°C. *k* -pH relation is expressed as linear equation of the form. -log(*k*) = *a* x pH + *C* and *t* _*1/2*_ -pH relation is expressed as linear equation of the form log(*t* _*1/2*_) = *b* x pH + *D*. Both are valid for the pH range from 6.0 to 10.0. R^2^ expresses the goodness of fit of the linear regression.

| AHL | Hydrolysis rate $k$ | | | Half-life time $t_{1/2}$ | | |
| --- | --- | --- | --- | --- | --- | --- |
| | $a$ | $C$ | $R^2$ | $b$ | $D$ | $R^2$ |
| C6 | -0.81 ± 0.07 | 9.0 ± 0.5 | 0.955 | -0.81 ± 0.07 | 8.9 ± 0.5 | 0.955 |
| C8 | -0.77 ± 0.06 | 8.9 ± 0.5 | 0.966 | -0.77 ± 0.05 | 8.7 ± 0.5 | 0.966 |

